# AAV-delivery of diacylglycerol kinase kappa achieves long-term rescue of *Fmr1*-KO mouse model deficits of fragile X syndrome

**DOI:** 10.1101/2021.04.14.439810

**Authors:** Karima Habbas, Oktay Cakil, Boglarka Zambo, Ricardos Tabet, Fabrice Riet, Doulaye Dembele, Jean-Louis Mandel, Michaël Hocquemiller, Ralph Laufer, Françoise Piguet, Hervé Moine

## Abstract

Fragile X syndrome (FXS) is the most frequent form of familial intellectual disability. It results from the lack of the RNA binding protein FMRP and is associated with the overactivation of signaling pathways downstream of mGluRI receptors and upstream of mRNA translation. We previously found that diacylglycerol kinase kappa (DGKk) is a main mRNA target of FMRP in cortical neurons. Here we show that diacylglycerol kinase kappa (DGKk), when modified as to become FMRP-independent and delivered into the brain of adolescent mice using adeno-associated viral vectors, corrects brain diacylglycerol and phosphatidic acid homeostasis and the main phenotypic behaviors of the Fmr1-KO mouse model of FXS. Thus, DGKk appears as a key triggering factor of FXS pathomechanism while providing a possible means of intervention for FXS gene therapy.

**One sentence summary:** *DGKk* gene therapy in *Fmr1*-KO mouse model

## INTRODUCTION

Fragile X syndrome (FXS) is a main cause of familial intellectual disability and autistic spectrum disorder (ASD) with a prevalence in general population estimated as 1 in 5000 males and 1 in 8000 females (Hagerman et al., 2017; Kaufmann et al., 2004). FXS is also generally associated with variable behavioral symptoms that can include anxiety, hyperactivity, hypersensitivity, stereotypies, memory deficits and sleeping problems. FXS results from the loss of the fragile X mental retardation protein (FMRP), an RNA binding protein associated to mRNAs and the translation machinery and whose absence in *Fmr1*-deleted mice (*Fmr1*-KO) recapitulates FXS-like phenotypes (1994; Mientjes et al., 2006), with perturbation of neuronal protein translation in hippocampus and cortex (Dolen et al., 2007; Qin et al., 2005). FMRP loss leads to mRNA translation increase of neuronal proteins resulting from an overactivation of metabotropic group 1 glutamate receptor (mGluR)-dependent local mRNA translation (Bear et al., 2004), including phosphatidylinositol 3-kinase enhancer (PIKE) (Gross et al., 2015), matrix metalloproteinase 9 (MMP9) (Gkogkas et al., 2014; Sidhu et al., 2014), glycogen synthase kinase 3 (GSK3) (Guo et al., 2012) and amyloid-β A4 protein (APP) (Pasciuto et al., 2015; Westmark et al., 2011). The mGluR overactivation is one well established triggering factor of FXS pathomechanism, and we recently showed that mGluR diacylglycerol (DAG) and phosphatidic (PA) dependent signaling upstream of local mRNA translation is disrupted by FMRP loss (Tabet et al., 2016a). The diacylglycerol kinase kappa (DGKk) transcript was identified in mouse cortical neurons as the mRNA species most bound by FMRP and with highest in vitro binding affinity. DGKk expression was found severely reduced in the brain of *Fmr1*-KO mouse and a perturbation of DAG/PA acid homeostasis was observed in *Fmr1*-KO cortical neurons and in the brain of FXS individuals, suggestive of a decreased DGK activity and altered DAG/PA signaling (Tabet et al., 2016a). DGKk knockdown in wild-type mouse brains recapitulated FXS-like behaviors, and overexpression of DGKk in *Fmr1*-KO hippocampal slices rescued their abnormal dendritic spine morphology (Tabet et al., 2016a). Overall, DGKk appears to play a key role in dendritic spine morphology and function and in the determination of FXS-like behaviors. DGKk loss of function was proposed to be at the origin of the various forms of abnormal synaptic signaling in FXS by causing altered DAG/PA signaling (Tabet et al., 2016b). Thus, being the most proximal downstream mediator of FMRP action, DGKk could represent an interesting actionable therapeutic target.

Here we show that DGKk expression is lost in FXS patients’ postmortem brains as previously shown in *Fmr1*-KO mice (Tabet et al., 2016a). We show that the N-terminal region of DGKk is important for its positive translational control by FMRP and the deletion of this region renders the protein (ΔN-DGKk) independent of FMRP, suggesting that FMRP alleviates a translation blockade within the N-terminal region. ΔN-DGKk is able to modulate protein synthesis rate and eIF4E phosphorylation in neurons. Moreover, ΔN-DGKk expression in *Fmr1*-KO mouse brain using adeno-associated virus Rh10 (AAVRh10) is able to correct brain lipid profile dysregulations and achieve long term behavioral rescue (over 8 weeks after injections) of *Fmr1*-KO mouse, providing a first proof of principle of *DGKk* gene therapy in a mouse model of FXS.

## RESULTS

### DGKk mRNA translation requires FMRP and DGKk N-terminal truncation alleviates FMRP control

We previously showed that the expression of diacylglycerol kinase kappa (DGKk) is strongly reduced in brain of *Fmr1*-KO mouse (Tabet et al., 2016a) in agreement with the fact that DGKk mRNA was identified as the most abundant mRNA species associated with FMRP by crosslinking immunoprecipitation in cortical neurons, and with the highest in vitro binding affinity. Like in *Fmr1*-KO mouse brain, DGKK expression is also lost in FXS post-mortem cerebellum compared to unaffected controls (**Fig. 1A**). These data suggest that DGKk requires FMRP for its proper expression. DGKk is mostly expressed in neurons and is almost absent in non-neuronal cells (Tabet et al., 2016a). We then tested if it is possible to recapitulate FMRP control in a non-neuronal cell system. We analyzed the influence of FMRP on the expression of HA-tagged mouse and human DGKk (**Fig. 1B, Sup Fig. 1A**) transfected into two different cell lines, Cos-1 and Hela. Knock-down of endogenous FMRP with siRNA severely reduced the expression level of mouse DGKk (mDGKk) compared to control siRNA treated Cos-1 cells (**Fig. 1BC**) indicating a strong FMRP requirement for DGKk expression. Human DGKk (hDGKk) level is also affected by FMRP knock-down, indicating that FMRP control is conserved on RNA sequences from different species (**Sup Fig. 1A**). mDGKk mRNA level is not influenced by the lack of FMRP (**Sup Fig. 1B**), while the protein level is severely reduced, supporting a control mechanism at the mRNA translational level. This is in agreement with our previous data in mouse brain indicating that DGKk transcript level is not affected by the loss of FMRP, and DGKk mRNA is less associated with polyribosomes (Tabet et al., 2016a). Noticeably, DGKk is the only DGK isozyme interacting strongly with FMRP (Tabet et al., 2016a) and bearing a long N-terminal extension constituted of unique proline-rich and EPAP repeated motives (Imai et al., 2005) (**Fig. 1B**). The N-terminal part of the protein might play a critical role in DGKk expression considering the repetitive nature of the EPAP domain at the beginning of the coding sequence. Thus, we generated a mDGKk construct lacking the first 696 bases following the start codon, encompassing the EPAP domain (ΔN-DGKk), for expression assessment in cells. ΔN-DGKk lead to a strong increase of expression (about ten-fold higher) (**Fig. 1D**). FMRP reduction did not affect ΔN-DGKk (**Fig. 1D**), suggesting that the N-terminal domain of DGKk is required for FMRP control and that ΔN-DGKk expression does no longer depend on the presence of FMRP. Therefore, ΔN-DGKk represents a potential therapeutic target for FXS gene therapy by bypassing the need for FMRP.

**Fig. 1:**
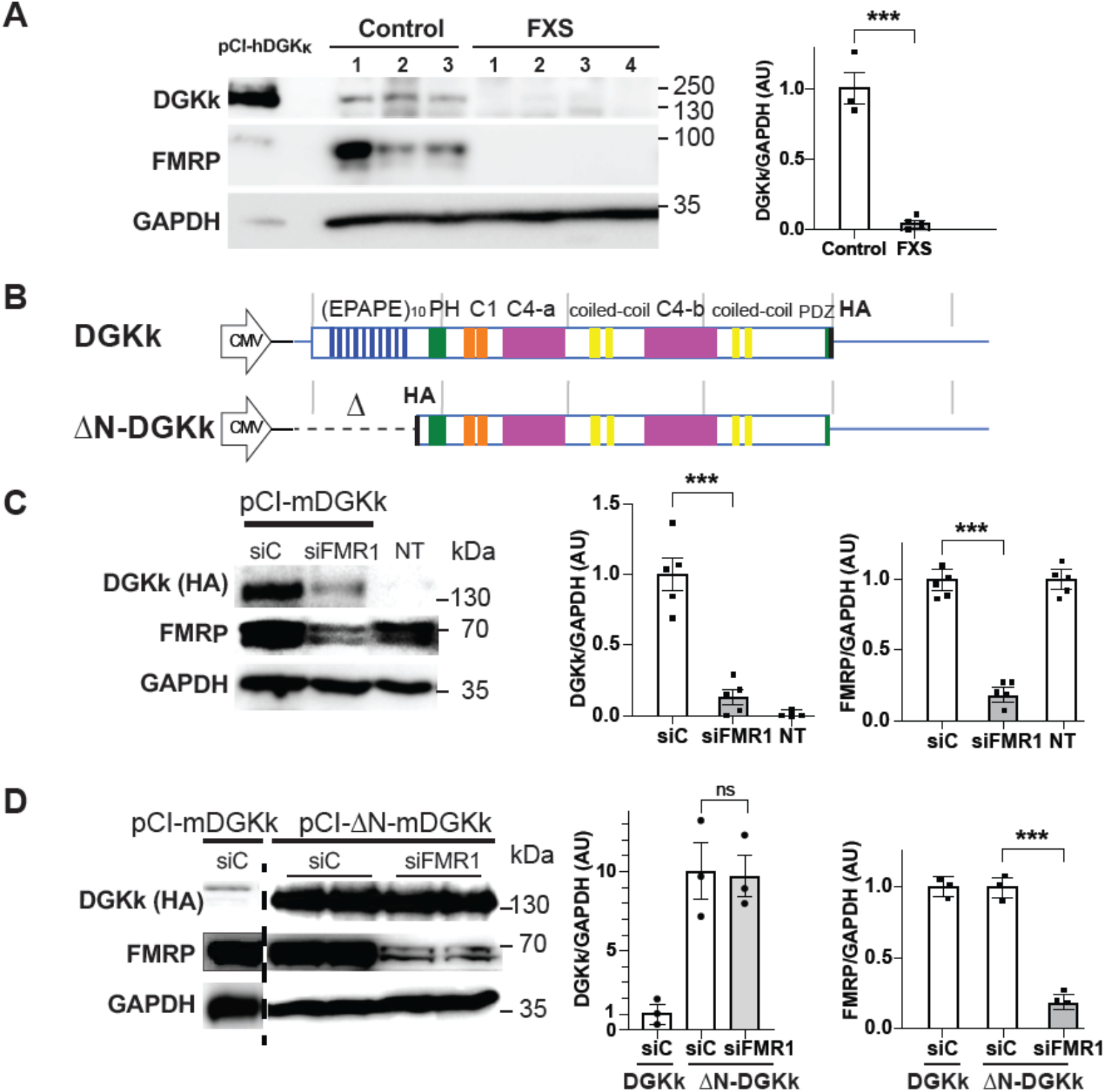
DGKk expression is altered in FXS and FMRP is required for translation control via its N-terminal domain. **A)** Western blot analysis of lysates from cerebellum of control (n = 3) and FXS patients ‘‘FXS’’ (n = 4). Hela cell extract transfected with pCI-hDGKκ was used as antibody specificity control and size marker. Representative images of immunoblots probed with antibodies against the indicated proteins are shown. GAPDH was the loading control. Quantification of Western blots is shown on right. Protein amounts of DGKk are normalized to GAPDH and presented as fold change relative to control. **B)** Schematic map of DGKk constructs used for the transfection experiments and subsequent vector preparations. The different domains of the protein are indicated and represented at scale (repeated EPAPE, Pleckstrin Homology PH domain, phorbol ester/diacyl glycerol binding C1 domain, catalytic split C4 a and b domains, putative PDZ binding motive, HA-tag, grey bars interval 1kB), 5’ and 3’ UTR regions are represented with blue line, 3’UTR not at scale (3.8 kB). **C)** Immunoblots and quantification of lysates from Cos-1 cells transfected with plasmid pCI-mDGKκ-HA or mock transfected (NT) and pre-transfected 24h before with siRNA control (siC) or against FMRP (siFMR1). GAPDH was used as a loading control. For quantification, the DGKk and FMRP signals were normalized against GAPDH signal and presented relative to the signal for siC treated cells (*n* = 5 in each group). **D)** Immunoblots and quantification of lysates from Cos-1 cells transfected with plasmid pCI-HA-ΔN-DGKκ or pCI-mDGKκ-HA and pre-treated with siRNA control (siC) or against FMRP (siFMR1). Quantifications as in C. Each point represents data from an individual culture, and all values are shown as mean ± SEM \*\*\**P* < 0.001, \*\**P* < 0.01, \**P* < 0.05 calculated by unpaired Student T test.

### ΔN-DGKk counteracts with FXS-like molecular defects

DGKk deregulation has been proposed to play a key role in FXS pathomechanism by altering DAG/PA signaling and leading to an excess of DAG and lack of PA potentially responsible of excessive protein synthesis, the major molecular hallmark of FMRP defects (Tabet et al., 2016a; Tabet et al., 2016b). To test the potential rescuing activity of ΔN-DGKk, we first analyzed its ability to modulate protein synthesis rate. Expression of ΔN-DGKk in Cos-1 cells reduced in a concentration dependent manner the protein translation rate, and this effect was counteracted by pretreatment of cells with DGK specific inhibitors R59022 and R59949 (Jiang et al., 2000) (**Fig. 2A**). These data demonstrate that ΔN-DGKk activity is upstream of mRNA translation and relies on conversion of DAG into PA, as DGK inhibitors prevent ΔN-mDGKk from reducing mRNA translation by blocking its enzymatic activity.

**Fig. 2:**
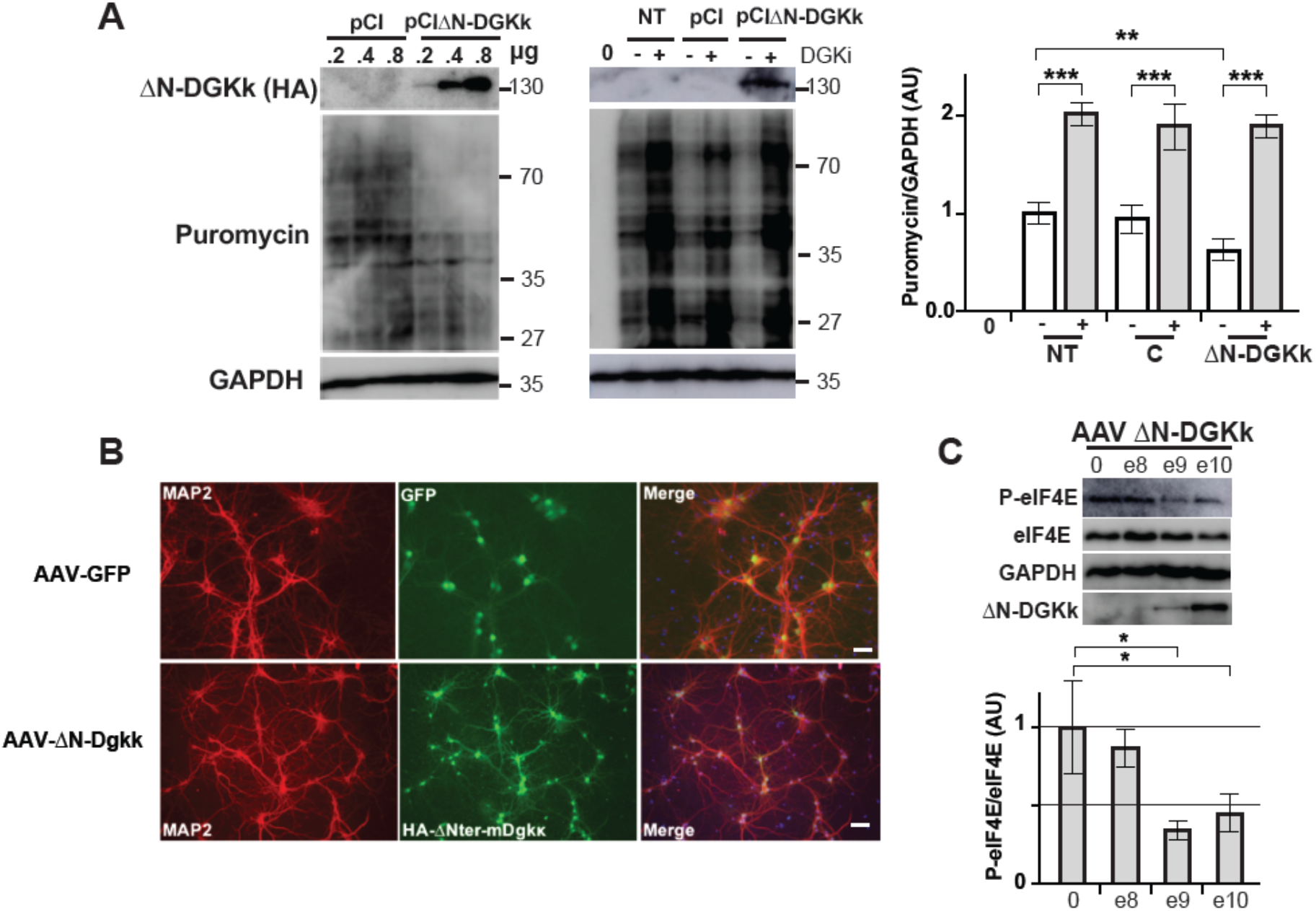
ΔN-DGKk expression impacts cellular signaling. **A)** Immunoblots and quantification of lysates from Cos-1 cells transfected with plasmid pCI-HA-ΔN-DGKκ, untransfected (NT), or plasmid pCI control, with the indicated amount of plasmid (μg) and incubated with puromycin to measure basal rates of protein synthesis. GAPDH was used as a loading control. 0 indicates no puromycin treatment, −/+ indicates treatment without or with DGK inhibitor (DGKi) 3 μM R59022 and 0.2 μM R59949 at 6 μM, 15 min. Densitogram of puromycin incorporation is presented as change relative to mock transfected conditions (*n* = 3). **B)** Representative immunofluorescence staining of cortical neuron cultures transduced at 8 DIV (days in vitro) with 10e9 VG/ml culture volume AAVRh10 GFP or ΔN-DGKk and assessed after 5 days using anti-MAP2 and anti-HA for ΔN-DGKk or direct 488 nm excitation for GFP, Dapi was used to visualize nuclei on merged images. Scale bar, 40 μm. **C)** Representative immunoblots of lysates from cortical neurons transduced with AAVRh10-ΔN-DGKk (AAV ΔN-DGKk), at the indicated titers (VG/ml culture volume) and quantification of phosphorylation and total levels of eIF4E. GAPDH was used as a loading control. For quantification, the phospho-protein signal was normalized against total protein signal and is presented relative to the signal for vehicle-treated cultures. Each point represents data from an individual culture, and all values are shown as mean ± SEM \**P* < 0.05 calculated by One way-ANOVA with Tukey’s multiple comparison test (*n* = 3 individual cultures).

To assess the biodistribution and efficacy of ΔN-DGKk in a FXS mouse model, we built our expression cassette to be driven by the neuron specific promoter synapsin and packaged into two different adeno-associated viral vectors (AAV), Rh10 and PHP.eB. Both AAV vectors harboring ΔN-DGKk demonstrated efficient neuronal transduction in *Fmr1*-KO cortical neurons, confirming that ΔN-DGKk can be expressed in the absence of FMRP (**Fig. 2B**). Phosphorylation of initiation factor eIF4E, which regulates protein synthesis by reducing translation initiation level (Sonenberg, 1994) and is increased after DAG-signaling activation (Wang et al., 1998), is increased in *Fmr1*-KO mouse and FXS patients (Gantois et al., 2017). We show that P-eIF4E is also increased in *Fmr1*-KO mouse cortical neurons compared to WT littermates (**Sup Fig. 2A**) and ΔN-DGKk is able to reduce their eIF4E phosphorylation (**Fig. 2C**). No sign of ΔN-DGKk-specific neuronal toxicity was observed in neuronal cultures transduced with AAV expressing ΔN-DGKk at high multiplicity of infection (MOI) (**Sup Fig. 2B-E**). An overall reduction in NeuN positive cells was observed upon treatment of cells with AAV, independently from ΔN-DGKk expression, as the same effect was observed with AAV-FMRP. Such effect was visible with a 10-fold lower amount of AAV-FMRP (10e9 VG/well) (**Sup Fig. 2D**). Additional tests (caspase 3/7, LDH) did not show signs of toxicity in vitro (**Sup Fig. 2B, C**).

### In vivo correction of phosphatidic acid level in adult mice using multiple routes of administration

ΔN-DGKk was administered to 5-week-old *Fmr1*-KO mice by intravenous injection of AAVPHP.eB-ΔN-DGKk or intracerebral injection into the striatum and hippocampus of AAVRh10-ΔN-DGKk, respectively. Single retro-orbital injection of AAVPHP.eB-ΔN-DGKk at 10^11 VG/mouse enabled 0.5-1 VG/cell throughout the brain (**Fig. 3AB**) four or eight weeks after injection, with low protein expression as visualized by Western blot (**Fig. 3C**) and immunohistochemistry (**Fig. 3D, Sup Fig. 3A**). Intracerebral injection of AAVRh10-ΔN-DGKk at 5×10^11 VG/mouse enabled higher brain transduction, with about 100 VG/cell in the hippocampus and about 25 VG/cell in rest of brain leading to high protein expression levels (**Fig. 3BCD**). AAVRh10 vector lead to robust ΔN-DGKk expression throughout the hippocampus (CA1 and CA2 regions mainly), cortex and striatum (**Fig. 3D, Sup Fig. 3A**). DGKk deregulation was shown to alter DAG/PA balance in neuronal cultures and human postmortem brains (Tabet et al., 2016a). We confirmed a marked decrease of total PA level (22% ±8) in the cortex of *Fmr1*-KO mice at 13 weeks of age (**Fig. 3E**). The other mouse brain regions did not show significant differences (**Sup Fig. 3B**). The decrease in the total pool of PA is reflected by a reduction of most PA species (including abundant species 34:1, 36:2, 38:4 or low species 38:3, 40:1) (**Sup Fig. 3D-F**). Thus, DGKk loss in *Fmr1*-KO cortex led to alteration of PA phosphorylation independent from fatty acid patterns. ΔN-DGKk delivery to the brain alleviated PA reduction in *Fmr1*-KO and led to a PA level comparable to vehicle injected WT mice (**Fig. 3E**). This is true for all fatty acid PAs with levels undistinguishable from control, demonstrating a complete rescue of PA by ΔN-DGKk expression (**Sup Fig. 3D-F**). Intravenous delivery of PHP.eB-ΔN-DGKk led to partial correction of total PA level (**Fig. 3E**) that was further confirmed at the single PA level, indicating that ΔN-DGKk is able to balance the PA with only few neurons transduced (**Sup Fig. 3D-F**). Total DAG level was not significantly altered in cortex (**Fig. 3F**) and other brain areas tested (**Sup Fig. 3C**), and at the level of individual DAG species (**Sup Fig. 3 G-I**). Nine other lipid classes tested did not show significant differences between the groups (**Sup Fig. 7J**) indicating that the effect of ΔN-DGKk is restricted to the correction of DAG/PA balance.

**Fig. 3:**
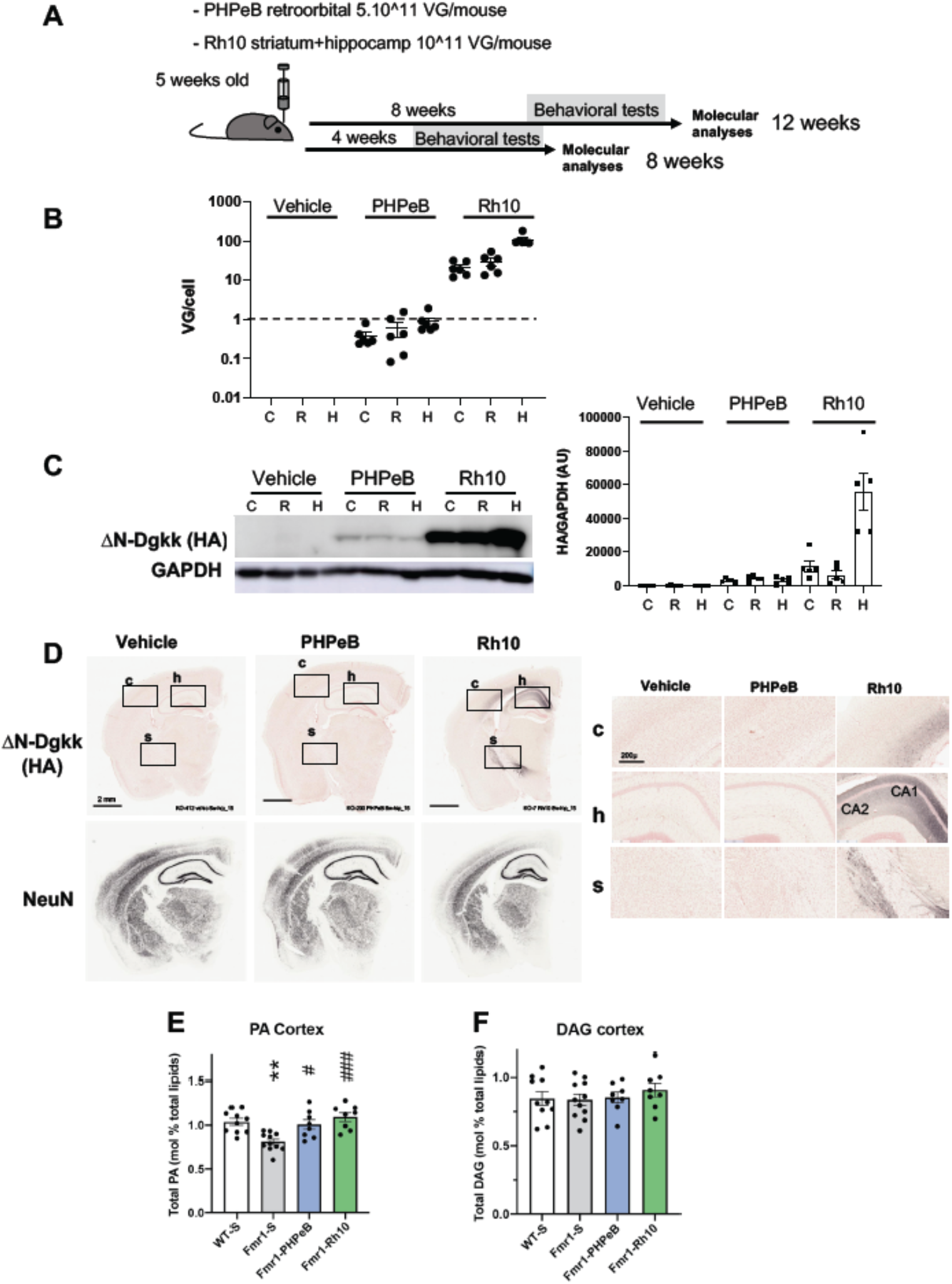
ΔN-DGKk expression in vivo with AVV vectors corrects abnormal cortical phosphatidic acid level of Fmr1-KO mice. **A)** Scheme of timeline analyses performed. **B)** Viral titers (viral genome copy VG per cell) determined by qPCR in cortical, hippocampal and rest of brain areas of Fmr1-KO mice treated with saline solution (Vehicle), AAVPHP.eB-ΔN-DGKk (PHP.eB), AAVRh10-ΔN-DGKk (Rh10) 8 weeks after injections. Data are mean ± SEM. Each dot represents an individual mouse. **C)** Immunoblots and quantification of ΔN-DGKk protein in lysates from brain areas of mice treated as in B. GAPDH was used as a loading control. Densitogram of ΔN-DGKk expression is presented as change relative to vehicle conditions. Data are mean ± SEM (*n* = 5). **D)** Representative coronal brain sections processed for detection of ΔN-DGKk using immunohistochemistry on Fmr1-KO mice treated with indicated treatment, 8 weeks post-injections, counter stained with eosin hematoxylin. Adjacent sections were immunolabelled with NeuN. 3 mice per genotype were processed. The sections shown are between Bregma levels –1.50 mm and –1.80 mm. Scale bar is 2 mm. Magnifications of regions of cortex (c), hippocampus (h), and striatum (s) are shown in side panels, scale bar 200 μm. **E)** Total phosphatidic acid (PA) level measure by mass spectrometry in cortex of WT mice treated with saline solution (WT-S) and Fmr1-KO mice treated with saline (Fmr1-S), AAVPHP.eB-ΔN-DGKk (Fmr1-PHP.eB), AAVRh10-ΔN-DGKk (Fmr1-Rh10) 8 weeks after injections. Data are expressed as mean ± SEM of mol % of total lipids and analyzed using one-way ANOVA and Tukey’s multiple comparisons test, **p<0.01, n=8 individual animals, except for WT n=7. **F)** Total diacylglycerol (DAG) level in cortex measured as in E).

### AAVRh10-ΔN-DGKk achieves long-term rescue of *Fmr1*-KO behavioral defects

Four weeks after injections, a battery of behavioral tests was performed on the AAV injected *Fmr1*-KO mice and their vehicle injected WT (WT-S) or *Fmr1*-KO (Fmr1-S) littermate controls (**Sup table 1**). Vehicle injected *Fmr1*-KO mice showed an increase of time spent and in number of entries in open arms of the elevated plus maze (EPM) compared to the WT-vehicle mice (1.9 and 1.8-fold of mean increase, respectively), suggesting decreased anxiety induced by the genotype. In contrast, AAVRh10-ΔN-DGKk treated mice (Fmr1-Rh10) showed no difference compared to WT-vehicle mice (**Fig. 4A**). A similar phenotype was observed in the open field arena of novel object recognition test (NOR), where vehicle injected *Fmr1*-KO mice showed an increase of time spent in the center of the arena, while AAVRh10-ΔN-DGKk treated mice were not different from WT-vehicle (**Fig. 4B**).

**Fig. 4:**
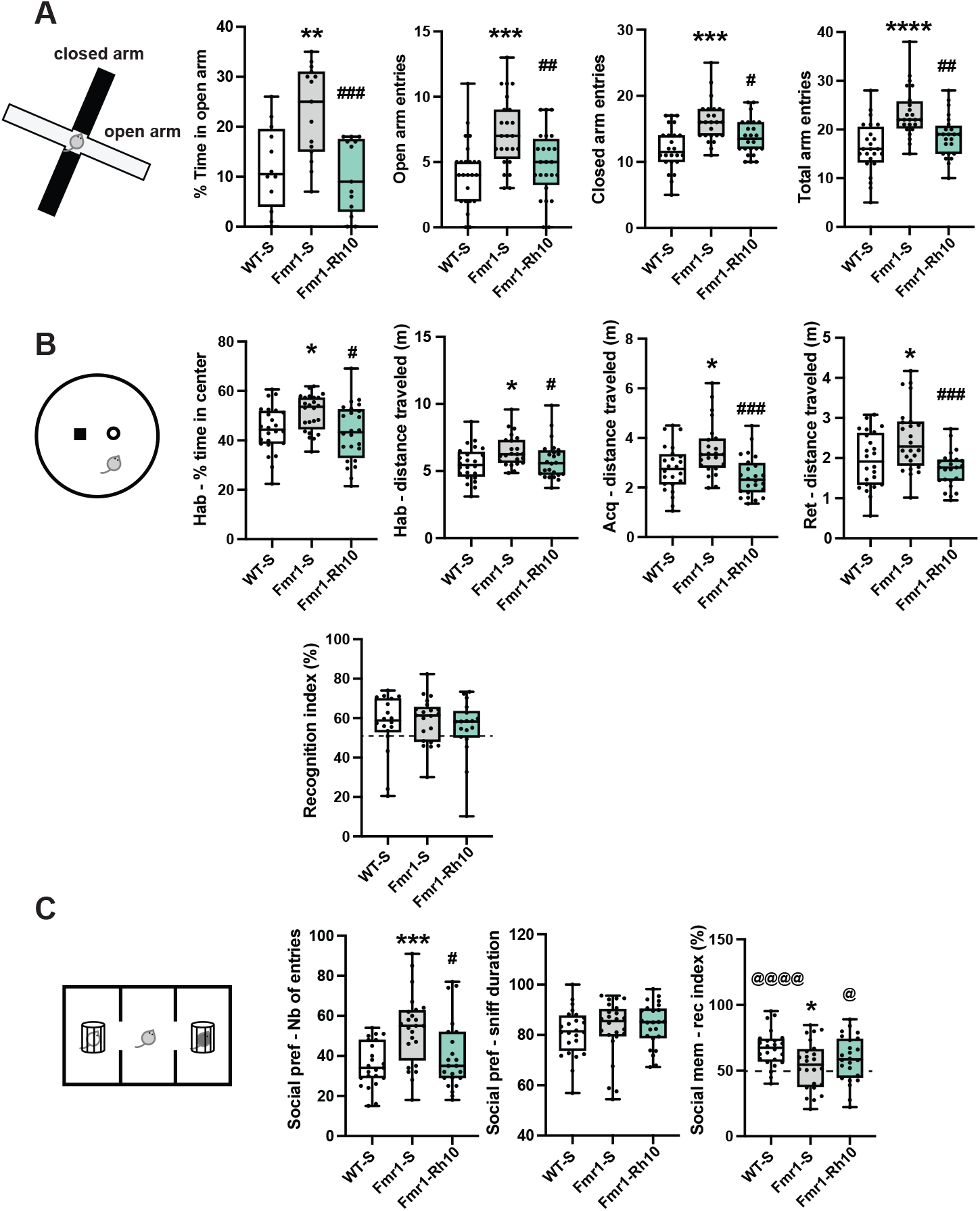
AAVRh10-ΔN-DGKk rescues behavior alterations of *Fmr1*-KO mouse 4 weeks after injections. **A)** Elevated Plus Maze. Percentage of time spent in open arms and number of entries in open, closed and total (open+closed) arms. **B)** Novel object recognition. Locomotor activity (distance in m) in the whole arena and percentage of time spent in the center during 15 min habituation. Locomotor activity during the acquisition and retention trials. Recognition index. **C)** Stereotypies. Number of digging. **D)** Social recognition. Number of entries in the two side compartments during preference session (left), social preference (percentage of exploration of a congener vs an object) (center) and social memory (percentage of exploration of a novel vs familiar congener) (right). Data are expressed as median with interquartile range with minimum and maximum values and analyzed using one-way ANOVA and Tukey’s multiple comparisons test and one group t-test. * p<0.05, **p<0.01, ***p<0.001, ****p<0.0001 vs WT-S; # p<0.05, ## p<0.01, ### p<0.001 vs Fmr1-S; @ p<0.05, @@ p<0.01, @@@@ p<0.0001 vs chance (50%).

Vehicle injected *Fmr1*-KO mice showed increased locomotor activity compared to WT-vehicle mice. This phenotype was seen in the EPM test (**Fig. 4A**, number of entries in open and closed arms), in the open field arena of NOR (**Fig. 4B**, distance travelled in habituation, acquisition and retention phases) and in the habituation phase of three-chambers social recognition test (**Fig. 4C**, number of entries), and was corrected in all tests with AAVRh10-ΔN-DGKk treatment. Signs of hyperactivity of *Fmr1*-KO mice were also seen in the first hour of light and dark phases of actimetry test (**Sup Fig. 4AB**), suggesting this phenotype is related to novelty of environment. AAVRh10-ΔN-DGKk treated mice did not show this phenotype. No difference in recognition index was observed between genotypes in the NOR test (**Fig. 4B**). Performing the NOR test in a smaller size arena ameliorated the recognition index of objects (**Sup Fig. 4D**), in correlation with longer object exploration times of objects (**Sup Fig. 4CD,** explorations). However, no difference was observed between the genotypes, suggesting an apparent lack of significant memory impairment of the *Fmr1*-KO mice. In small size arena the hyperlocomotor activity phenotype of *Fmr1*-KO mice was not visible (**Sup Fig. 4D**), potentially because this environment was less anxiogenic than a larger arena. *Fmr1*-KO vehicle mice showed a trend to dig more, to bury more marbles and to spend less time in grooming compared to the WT-vehicle mice, but these differences were not statistically significant (**Sup Fig. 4E**). ΔN-DGKk treatment seemed to counteract these effects, but the high variability of these phenotypes between mice precluded drawing conclusions.

Vehicle injected *Fmr1*-KO mice showed an increased number of entries in the three-chambers social recognition test and this phenotype was corrected with AAVRh10-ΔN-DGKk (**Fig. 4C, Sup Fig. 4F**). For social preference over object, no difference was observed between genotype groups (**Fig. 4C, Sup Fig. 4F**) indicating there was no significant effect of mutation or treatment on social preference. In the social memory test, *Fmr1*-KO mice failed to recognize previously encountered mice, while *Fmr1*-KO mice treated with AAVRh10-ΔN-DGKk had a higher recognition index relative to untreated mice (**Fig. 4C**).

Vehicle injected *Fmr1*-KO mice also showed a reduced ability to build a nest after 5 and 24h. AAVRh10-ΔN-DGKk treatment rescued this phenotype (**Sup Fig. 4G**).

The phenotypes observed at 4 weeks after injections were generally recapitulated in older animals at 8 weeks after injection (**Sup Fig. 5A-F)**. Consistent with the stability of the treatment effect, no significant variation of ΔN-DGKk protein level was observed between 8 and 12 weeks after AAV injection (corresponding to animal groups phenotyped at 4 and 8 weeks, respectively) (**Sup Fig. 3A**). *Fmr1*-KO-vehicle mice tested at 8 weeks showed reduced recognition index in NOR test compared to WT-vehicle and *Fmr1*-KO-Rh10-ΔN-DGKk (**Sup Fig. 5D**), but within a smaller test arena where *Fmr1*-KO-vehicle mice exhibit less hyperactivity (**Sup Fig. 5E**) all groups performed equally, suggesting that the memory performance alteration of *Fmr1*-KO mice might be caused by hyperactivity. *Fmr1*-KO-vehicle mice tested at 8 weeks after injection (13-week-old) showed a strong digging behavior compared to WT-vehicle and *Fmr1*-KO-Rh10-ΔN-DGKk, a phenotype that was not visible at 4 weeks (**Sup Fig. 5F**). An increase in body weight was observed in *Fmr1*-KO-vehicle group compared to WT-vehicle. WT-vehicle and *Fmr1*-KO-Rh10-ΔN-DGKk showed no weight difference, suggesting the treatment rescued this phenotype (**Sup Fig. 5G**). *Fmr1*-KO-vehicle phenotypes were reproduced in another cohort aimed at testing retro-orbital administration of PHP.eB-ΔN-DGKk (**Sup Fig. 6A-F**). But unlike the mice which received AAVRh10-ΔN-DGKk, (and except for distance travelled during retention phase of NOR, **Sup Fig. 6D)**, *Fmr1*-KO-PHP.eB-ΔN-DGKk mice showed no significant improvement compared to *Fmr1*-KO-vehicle, suggesting that the level of ΔN-DGKk, although well distributed throughout the brain, was too low to achieve a sufficient effect. Macroorchidism, a well-established phenotype of *Fmr1*-KO model, was not found corrected by the treatments (**Sup Fig. 7**), possibly reflecting a non-neuronal origin.

## DISCUSSION

FXS is currently uncured as no disease-modifying treatment could be validated despite several clinical trials with investigational drugs (Yamasue et al., 2019). Our data provide evidence that neuron targeted expression of DGKk enzyme with an AAV-based gene therapy approach is able to provide long-term correction of the main behavioral deficits in the young adult *Fmr1*-KO mouse model of FXS (**Sup table 2**).

DGKk is an enzyme whose mRNA was previously found to be the main target of FMRP in cortical neurons (Tabet et al., 2016a). We show that DGKk expression is strongly dependent on FMRP and severely altered in FXS brain. In fact, no other protein has been demonstrated to be so critically dependent upon FMRP. Loss of DGKk activity could play a critical role in manifestation of FXS phenotypes because it is a master regulator of second messenger lipids DAG/PA balance and has the potential to be the triggering cause of the many altered neuronal signaling pathways observed in FXS (Tabet et al., 2016b). Removal of the N-terminal part of DGKk does not impact DGKk activity in vitro (Imai et al., 2005), while it abolishes regulation by FMRP. The FMRP-independent ΔN-DGKk protein conserved its ability to modulate cell signaling and showed capacity to rescue a fully developed *Fmr1*-KO mouse brain. At the dose used, intravenous administration of AAVPHP.eB-ΔN-DGKk was unable to rescue the *Fmr1*-KO phenotype, presumably because of insufficient, albeit homogenous expression in the brain. In contrast, intracerebral injection of AAVRh10-ΔN-DGKk led to higher expression of ΔN-DGKk in the mouse brain and correction of disease phenotypes, without affecting survival and with no signs of toxicity several weeks after dosing.

Consistently, high ΔN-DGKk expression in neuronal cultures was not associated with cellular toxicity. Rescuing *Fmr1*-KO mouse with AAV-based FMRP administration at an early stage (P0-P5) has provided the first proof of concept of a gene therapy approach for FXS (Gholizadeh et al., 2014; Hampson et al., 2019), but also revealed that inappropriate levels of FMRP expression can lead to worsening of FXS phenotypes, possibly due to the fact that FMRP, like most RNA binding proteins, induces cellular stress when overexpressed (Mazroui et al., 2002). Overall, rescue of *Fmr1*-KO phenotypes with ΔN-DGKk expression in neurons strengthened the notion that DAG/PA imbalance in neurons is a critical factor of the disease and acting on this imbalance is beneficial for FXS-like condition, including at a late stage of development. Use of ΔN-DGKk could offer potential for FXS gene therapy, representing a very specific target in the complex pathomechanism of the disease.

## MATERIAL AND METHODS

### Ethics statement

Animal work involved in this study was conducted according to ARRIVE guidelines and received authorization from relevant national (Comité National de Réflexion Ethique en Expérimentation Animale).

### Cloning of human and mouse DGKκ mRNA

Mouse DGKκ was subcloned from clone IMAGE IRAVp968H03163D. The missing 3′ UTR and 5′ UTR-N-ter region were cloned by PCR from mouse genomic DNA with primer sets (GCAGCTAGCTCCTTGAAAGCTGGAAGGAGA and AATAGAATGCGGCC-GCCAGCTTCAACAGCACTTGTAG) and (CCAgtcgacTTAGACCTCAGAGCTGCGCTAGC and CCAgctagcCCAGGACTCTGGGGCCCTCTCCAT), respectively. The 3’ UTR region was introduced at XbaI and NotI sites of the pYX-ΔN DGKκ vector to give pYX-ΔN-DGKκ-3’UTR, and the 5’-UTR-Nter region at SalI-NheI sites of the pYX-DGKκ-3’UTR, NheI site was subsequently deleted by PCR mutagenesis. pCI-mDGKκ-HA and pCI-HA-ΔN-DGKκ were obtained by PCR subcloning into pCI vector (GenBank U47119) with addition of the HA sequence before the STOP codon or after ATG, respectively. Human hDGKκ was subcloned from plasmid pAcGFPC1humDGKk (Imai et al., 2005) into Nhe1 of pCI vector with addition of 5’ and 3’UTRs by PCR cloning to produce pCI-hDGKκ.

### AAV ΔN-DGKk vectors construction and preparation

ΔN-DGKk HA-tagged DGKκ was cloned under the control of the hSynapsin promoter replacing EGFP in the control plasmid pENN.AAV.hSynapsin.EGFP.RBG (provided by the Penn Vector Core at University of Pennsylvania, Philadelphia) to give pAAV-ΔN-DGKk. Recombinant adeno-associated virus serotype 9 (AAV9), Rh10 (AAVRh10), PHP.eB (AAVPHP.eB) production was carried out by using the AAV Helper-Free system (Agilent Technologies) with some modifications. AAV vectors were generated by triple transfection of 293T/17 cell line using Polyethylenimine (PEI) and plasmids pAAV-hsynapsin-HA-ΔN-DGKκ or pENN.AAV.hSynapsin.EGFP.RBG together with pHelper (Agilent) and pAAV2/9 or pAAV2/Rh10 (provided by J.Wilson and J.Johnston at Penn Vector Core), or pUCmini-iCAP-PHP.eB (provided by V.Gradinaru and J.Johnston) for serotypes 9, Rh10 and PHP.eB, respectively. Two days after transfection, cells were collected, lysed by three freeze/thaw cycles in dry ice-ethanol and 37 °C baths, further treated with 100 U/mL Benzonase (Novagen) for 30 min at 37 °C, and clarified by centrifugation at 3,000 × g for 15min. Viral vectors were purified by iodixanol (Optiprep, Axis Shield) gradient ultracentrifugation followed by dialysis and concentration against PBS containing 0.5 mM MgCl_2_ using centrifugal filters (Amicon Ultra-15.100 K) and filtered through 0.22u (Zolotukhin et al., 2002). Viral particles were quantified by real-time PCR Q-PCR using LightCycler480 SYBR Green I Master (Roche) and primers targeting the flanking sequence of ITR2 (GTAGATAAGTAGCATGGC and CTCCATCACTAGGGGTTCCTTG) or the flanking sequence of rabbit β-globin polyadenylation signal (CCCTTGAGCATCTGACTTCTGG and AGGGTAATGGGTATTATGGGTGGT). To achieve comparable working concentrations, viruses were diluted to a final concentration of 1x 10^13^ viral genome per ml (VG/ml) and stored at −80°C until use.

### Cell culture and transfections

COS-1 cells were grown in DMEM supplemented with 10% (v/v) FCS and 1 g/L glucose in the presence of antibiotics at 37 °C in 5% CO2. The day before transfection, 4×10^4^ cells were plated into 24-well format plates in 500 μL of antibiotic-free medium. Transfections of plasmids were performed in triplicate with Lipofectamine 2000 (Invitrogen) as directed by the manufacturer with 10 pmol of siRNA (Control or On Target plus Smart Pool mouse *FMR1*; Thermo Fisher Scientific) in a final volume of 600 μL. Twenty-four hours later, 300 ng of the reporter pCI-DGKκ-HA or pCI-HA-ΔN-DGKκ and 10 nM shRNA were cotransfected as above. Twenty-four hours later, cells washed twice in PBS were lysed directly in the loading buffer [100 mM Tris·HCl, pH 6.8, 4% (w/v) SDS, 30% (v/v) glycerol, 1.4M β-mercaptoethanol, and bromophenol blue] for 3 min at 95 °C.

### Primary cortical neuron cultures and treatments

Neuron cultures were performed as described in (Tabet et al. 2016). Briefly, cortices from C57BL/6J *Fmr1*+/y or *Fmr1*−/y mouse embryos (embryonic day E17.5) were dissected in 1xPBS, 2.56 mg/mL D-glucose, 3 mg/mL BSA, and 1.16 mM MgSO4, incubated for 20 min with 0.25 mg/mL trypsin and 0.08 mg/mL DNase I, and mechanically dissociated after supplementation of medium with 0.5 mg/mL trypsin soybean inhibitor, 0.08 mg of DNase I and 1.5 mM MgSO_4_. The cells were plated on poly-L-lysine hydrobromide-coated six-well culture plates for 8 days in Neurobasal Medium (GIBCO) supplemented with B27, penicillin/streptomycin, and 0.5 μM L-glutamine. Where indicated, cultures were treated with addition of puromycin solution at the indicated concentrations and times. DGK inhibitors R59022 (DGK Inhibitor I; Calbiochem) and R59949 (DGK Inhibitor II; Calbiochem) were applied at concentrations of 3 and 0.2 μM each for 15 min at 37 °C. After treatment, cells were immediately washed with ice-cold PBS and lysed in 4X Laemmli buffer.

### Western blot analyses

Immunoblotting was performed as described previously (Tabet et al., 2016a). Proteins (equivalent to 15 μg) were denatured 5 min at 95°C and resolved by 10% SDS-PAGE. Separated proteins were transferred onto PVDF Immobilon P membrane (Millipore) using a Mini Trans-Blot (Biorad) cell. Membranes were blocked for 1h with TBS-T 1X (Tris-Buffer Saline, pH 7.4 and 0.1% Tween-20 v/v) containing 5% (w/v) BSA or 5% nonfat dry milk. Membranes were incubated overnight at 4°C with primary antibodies diluted in TBS-T buffer containing 5% w/v BSA or milk as follows: mouse anti-FMRP (1C3, 1:10,000, IGBMC), purified mouse anti-HA.11 16B12 (1:5000, Biolegend), rabbit anti-hDGKk (1:1000, PA5-25046, ThermoFisher), anti-p-EIF4e Ser209 (1:1000, #9741 Cell Signaling), anti-EIF4 (1:1000 BSA, #9742, Cell Signaling), anti-puromycin (12D10, 1:2000, Sigma-Aldrich), anti-S6 (5G10, 1:1000 #2217S, Cell Signaling), anti-p-S6 (Ser235/236, 1:1000, 2211S Cell Signaling), anti-GAPDH (MAB374, 1:10.000, Merck) was used as an internal standard. Membranes were washed in TBS-T buffer and then incubated for an hour at room temperature with the corresponding horseradish peroxidase-conjugated pre-adsorbed secondary antibody (1:5000, blocking solution corresponding, Molecular Probes). Membranes were washed in TBS-T buffer and immunoreactive bands were visualized with the SuperSignal West Pico Chemiluminescent Substrate (Pierce). Immunoblot pictures were acquired using LAS600 GE Amersham and density of the resulting bands was quantified using ImageJ and statistical significance assessed using repeated measures analysis of variance (ANOVA) with Fisher’s post hoc comparisons.

### Analysis of cortical neuron cultures by immunofluorescence microscopy

Primary cortical neurons grown on poly-L-lysine hydrobromide coated glass coverslips were fixed in 4% (w/v) paraformaldehyde (PFA) in 1xPBS at room temperature (RT) for 20 min, permeabilized with 1X PBS and 0.2% Triton X-100 for 10 min at RT, and blocked for 1 h in 1x PBS and 0.1% Triton X-100 with 5% (w/v) BSA. Neurons were incubated with primary antibodies rabbit anti-MAP2 AB5622 (1:500, Merck Millipore), mouse anti-HA.11 16B12 (1:500, Biolegend), mouse anti-NeuN (1:500, MAB377 Merck), rabbit anti-GFAP (1:500, 173002, Synaptic system), overnight at 4 °C. After three washes in 1X PBS and 0.1% Triton X-100 for 10 min, neurons were incubated with secondary goat antibody anti-rabbit (Alexa Fluor 594, 1:1.000, Invitrogen) and anti-mouse (Alexa Fluor 488, 1:1000, Invitrogen), for 1h at RT, and subsequently washed three times in 1xPBS and 0.1% Triton X-100 for 10 min. Coverslips were mounted with antifading medium (Vectashield, Vector) with DAPI and analyzed by fluorescence microscopy. Images were acquired with CellInsight CX7 (Thermo Scientific) using a 10x objective and analyzed with HCS Studio Cell Analysis Software (nuclear segmentation, NeuN and GFAP intensities). Quantification of positive cells for each of these staining was done by applying a threshold manually, based on nuclear segmentation and across 81 fields. Percentage of cells positive for NeuN and GFAP staining was quantified for each well.

### Caspase 3/7 activity detection

Caspase 3/7 positives cells were determined with CellEvent Caspase-3/7 Green Detection Reagent following manufacturer instructions. Briefly, primary neurons were pepared as for LDH assays, treated with reagent diluted at 8 μM in 5% FBS NBM. Positive control wells were treated with apoptotic inducer staurosporine at 0.1 and 1uM for 6 hours. Cells were fixed with were fixed in 4% (w/v) PFA and nuclei were counterstained with DAPI. Images were acquired with CellInsight CX7 (Thermo Scientific) using a 10x objective and analyzed with HCS Studio Cell Analysis Software (nuclear segmentation and casp3/7 intensities). Quantification of percentage of caspase-3/7 positive cells was done for each well by applying a threshold manually, based on nuclear segmentation and across 81 fields.

### Lactate dehydrogenase releasing assay

Lactate dehydrogenase (LDH) release was determined with the Cytotoxicity Detection KitPLUS kit (Roche) following manufacturer instructions. Briefly, primary neurons from Fmr1^+/y^ or Fmr1^−/y^ E17.5 embryos plated at 300,000 cells/well in 24-well plate and transduced at 7 DIV with indicated AAV were tested after 7 days. LDH release was measured in microplate reader at 490 nm after 2 hours at 37°C with reaction medium. Maximum LDH release was measured in same conditions after 30min at RT with stop solution. The % of cell death was determined using the formula: % cell death = experimental LDH release /maximum LDH release.

### Animal housing

At weaning age (4 weeks), animals were grouped by 3 or 4 individuals from same age and genotype in individually ventilated cages (GM500, Tecniplast, UK), with poplar shaving bedding (Lignocell Select, JRS, Germany), and maintained under standard conditions, on a 12-h light/dark cycle (7h/19h), with standard diet food (standard diet D04, Scientific Animal Food and Engineering, France) and water available ad libitum. Mice from a same cage received the same treatment and were transferred in the animal facility of the phenotyping area the next week.

### Stereotaxic Surgery and AAV Injections

Five-week old mice C57BL/6J *Fmr1*-/y or C57BL/6J *Fmr1*+/y littermates were deeply anesthetized with ketamine/xylazine (Virbac/Bayer, 100/10 mg/kg, 13 mL/kg, intraperitoneal) dissolved in sterile isotonic saline (NaCl 0.9%) and mounted onto a stereotaxic frame (World Precision Instruments). AAVRh10-DGKκ were injected bilaterally into the striatum (coordinates relative to bregma: anterior-posterior + 0.5 mm; lateral = ±2.2 mm; vertical −3.5 mm) and hippocampus (coordinates relative to bregma: anterior-posterior - 1.7 mm; lateral = ±1.5 mm; vertical −2.0 mm) according to the mouse brain atlas (Paxinos G, Franklin KBJ (2001)). A volume of 2.5 μL of AAV vector (corresponding to 10^e^11 Genome copies) or saline solution was delivered bilaterally per site of injection with a slow injection rate (0.2 μL/min) through a 32-gauge small hub removable needle mounted on a 10 μL Hamilton syringe connected to a micropump (World Precision Instruments). After each injection was completed, the injector was left in place for an additional 2 min to ensure optimal diffusion and minimize backflow while withdrawing the injector.

### Retroorbital AAV Injections

Five-week old mice C57BL/6J *Fmr1*-/y or C57BL/6J *Fmr1*+/y littermates were deeply anesthetized with ketamine/xylazine (Virbac/Bayer, 100/10 mg/kg, 10 mL/kg, intraperitoneal) dissolved in sterile isotonic saline (NaCl 0.9%). AAVPHP.eB-DGKκ were injected retroorbitaly with a volume of 80 μL.

### Analysis of ΔN-DGKk expression in brain sections

Freshly dissected brains were fixed overnight in PFA 4% and stored in PBS1X prior being processed following Neuroscience Associates procedure https://www.neuroscienceassociates.com/technologies/multibrain/. Half-brains were washed in PBS1X solution and embedded in gelatin matrix. MultiBrain^®^ cryosections were prepared with 30μ thickness for free-floating immunolabeling with anti-HA (3F10, 1:150, Sigma) and counter stained with hematoxylin/eosin. Adjacent sections were immunolabelled with anti-NeuN (1:150, MAB377 Merck).

### Behavioral Experiments

Behavioral experiments were conducted 4 weeks or 8 weeks after AAV injections to allow sufficient time for viral transduction and DGKκ expression. Effective gene expression was assessed by q-PCR to measure viral titer, by western blot and by immuno-histochemistry in 3 different brain areas (cortex, hippocampus and rest of brain). Phenotyping pipeline is described in table 1.

### Circadian Activity

Spontaneous locomotor activity and rears are measured using individual cages (20 × 10 × 8 cm) equipped with infra-red captors. The quantity of water and food consumed is measured during the test period using automated pellet feeder and lickometer (Imetronic, Pessac, France). Mice are tested for 32 hours in order to measure habituation to the apparatus as well as nocturnal and diurnal activities. Results are expressed per 1 h periods and/or as a total of the different activities.

### Elevated plus maze

The apparatus used is completely automated and made of PVC (Imetronic, Pessac, France). It consists of two open arms (30 × 5 cm) opposite one to the other and crossed by two enclosed arms (30 × 5 × 15 cm). The apparatus is equipped with infrared captors allowing the detection of the mouse in the enclosed arms and different areas of the open arms. Mice were tested for 5 min during which the number of entries into and time spent in the open arms were measured and used as an index of anxiety. Closed arm entries and total arm entries were used as measures of general motor activity.

### Novel object recognition task

Mice were tested in a circular arena (50cm diameter and 30cm height basin). The locomotor activity was recorded with the EthoVision XT video tracking system (Noldus, Wageningen, Netherlands). The arena was virtually divided into central and peripheral regions and homogeneously illuminated at 40 Lux. Animals were first habituated to the arena for 15 min. Each mouse was placed in the periphery of the arena and allowed to explore freely the apparatus, with the experimenter out of the animal’s sight. The distance traveled and time spent in the central and peripheral regions were recorded over the test session. The percentage of time spent in center area was used as index of emotionality/anxiety. The next day, mice were tested for object recognition in the same arena. They were submitted to a 10-minutes acquisition trial during which they were placed in the arena in presence of a sample objects (A and A’) (2.5 cm diameter marble or 2 cm edge plastic dice). The time the animal took to explore the samples (sniffing) was manually recorded. A 10-minutes retention trial was performed 24 h later. During this trial, one of the samples A and another object B (marble or dice depending on acquisition) were placed in the open-field, and the times tA and tB the animal took to explore the two objects were recorded. A recognition index (RI) was defined as (tB / (tA + tB)) ×100.

### Nest building

On the day of test, mice were singly transferred in a standard cage for the duration of nest building measurement. A block of nesting material (5×5cm hemp square, Happi Mats, Utopia) was placed in the cage. Pictures were taken and visual scoring occurred at 2, 5, 24 h without disturbing the animals. The room temperature was noted when the nest was scored, since nest building has a thermoregulatory function and therefore may be influenced by ambient temperatures. We used a 0-5 scale described by (Gaskill et al., 2013): 0 = undisturbed nesting material; 1 = disturbed nesting material but no nest site; 2 = a flat nest without walls; 3 = a cup nest with a wall less than ½ the height of a dome that would cover a mouse; 4 = an incomplete dome with a wall ½ the height of a dome; 5 = a complete dome with walls taller than ½ the height of a dome, which may or may not fully enclose the nest.

### Social recognition test

Social recognition test evaluates the preference of a mouse for a congener as compared to an object placed in an opposite compartment. This test is also used for evaluation of social memory by measuring exploration of a novel congener as compared to a familiar one. Social behavior is altered in several diseases such as autism and mental retardation. The apparatus is a transparent cage composed with a central starting compartment and 2 side compartments where circular grid cup (goal box) is placed at each extremity, and where the congener can be placed during testing. Testing was performed for 2 consecutive days. On the first day, the mouse was placed in central box then allowed to explore freely the apparatus for 10 min in order to attenuate their emotionality. On the second day, a C57Bl/6 congener from the same sex was placed in one goal box and an object was placed in the opposite one. The mouse was then placed in the starting central compartment and allowed to explore freely the apparatus for 10 min. The position of the congener and object boxes was counterbalanced to avoid any potential spatial preference. The duration of exploration of each goal box (when the mouse is sniffing the grid delimiting the goal box) was manually measured and the percentage of time the mouse took to explore the congener was used as index of social preference (recognition preference). A 10min retention trial was then performed during which the object was replaced by a novel congener. The duration of exploration of each goal box was manually measured and the percentage of time the mouse takes to explore the congener was used as index of social memory. The social preference index (SR) is defined as (time Congener / (time Object + time Congener)) ×100; and the social memory index as (time novel Congener / (familiar congener + time novel Congener)) ×100.

### Lipidomic analyzes

Nitrogen frozen brain samples (cortex, hippocampus, rest) were let thaw on ice and mechanically homogenized with 1 vol H_2_0 with Precellys 24 system during 2×15sec at 4°C and 5300 rpm. Protein concentration of sample was adjusted at 5 mg/ml concentration and lipids were analyzed on Lipotype GmbH platform. Lipids were extracted using chloroform and methanol (Sampaio et al., 2011) with Hamilton Robotics STARlet. Samples were spiked with lipid class-specific internal standards prior to extraction. After drying and resuspending in MS acquisition mixture, lipid extracts were subjected to mass spectrometric analysis. Mass spectra were acquired on a hybrid quadrupole/Orbitrap mass spectrometer (Thermo Scientific Q-Exactive) equipped with an automated nano-flow electrospray ion source in both positive and negative ion mode. Lipid identification using LipotypeXplorer (Herzog et al., 2011) was performed on unprocessed (*.raw format) mass spectra. For MS-only mode, lipid identification was based on the molecular masses of the intact molecules. MSMS mode included the collision induced fragmentation of lipid molecules and lipid identification was based on both the intact masses and the masses of the fragments. Prior to normalization and further statistical analysis lipid identifications were filtered according to mass accuracy, occupation threshold, noise and background. Intensity of lipid class-specific internal standards was used for lipid quantification. The identified lipid molecules were quantified by normalization to a lipid class specific internal standard. The amounts in pmol of individual lipid molecules (species of subspecies) of a given lipid class were summed to yield the total amount of the lipid class. The amounts of the lipid classes were normalized to the total lipid amount yielding mol% per total lipids.

### Statistical analyses

Quantitative data were analyzed using single or repeated measures analysis of variance (ANOVA) and Student Newman Keuls test. For the comparison with chance, one group t-test was used. Qualitative parameters (nesting) were analyzed using χ2 test. The level of significance was set at p < 0.05.

## Acknowledgements

We thank Flora Tassone and Veronica Martinez-Cerdeno for human brain materials. We are indebted to Antonina Fedorova, Hélène Puccio, Yann Herault, Sirine Souali-crespo, Tania Sorg-Guss and members of Lysogene team for discussions and suggestions. We are grateful to Andréa Geoffroy and members of Institut Clinique de la Souris for help with the behavioral studies, Pascale Koebel for AAV preparations, Eric Flatter for technical assistance, Nadine Banquart-Ott and Chadia Nahy for mice care, Anne Maglott for toxicity analyses, Erwan Grandgirard for help with imaging, IGBMC core facilities, Zaiane Schmitt and Fumi Hoshino for technical assistance. We specially thank Laura Benkemoun and Fondation Maladies Rares for support and suggestions. We are grateful to colleagues of J. Chelly team for discussions.

## Author contributions

K.H., O.C, B.Z., and F.P. designed and performed experiments and helped writing the manuscript. D.D. performed statistical analyses. R.T., M.H., R.L. helped designing the experiments and writing manuscript. H.M. supervised the project, designed experiments and wrote the manuscript.

## Supplementary material

**Supplementary Fig. 1:**
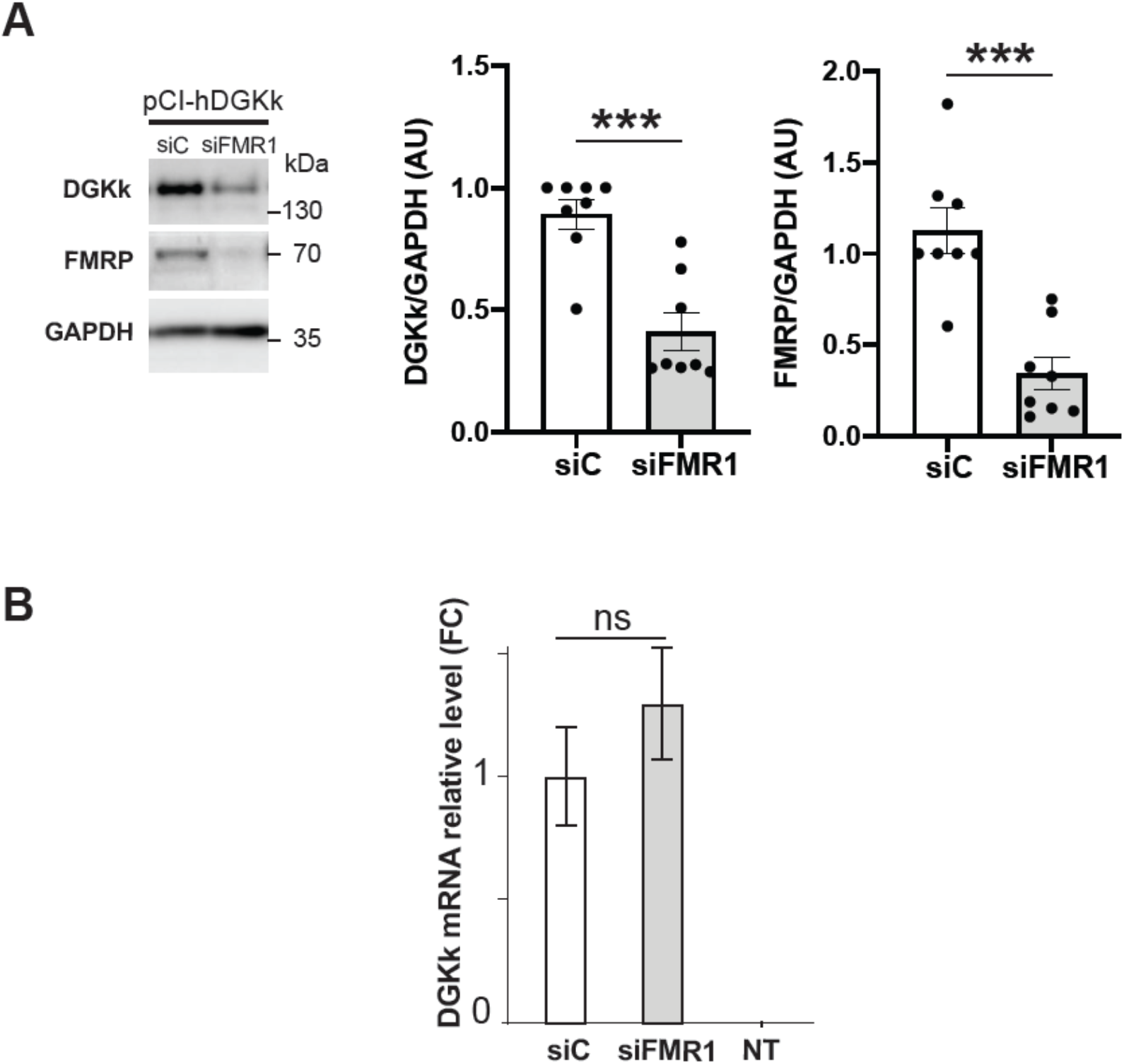
Influence of FMRP protein on human DGKk protein expression and on mouse DGKk mRNA level. **A)** Immunoblots and quantification of lysates from Hela cells transfected with plasmid pCI-hDGKκ encoding human DGKk and pre-transfected 24h before with siRNA control (siC) or against FMRP (siFMR1). GAPDH was used as a loading control. For quantification, DGKk and FMRP signals were normalized against GAPDH signal and presented relative to the signal for siC treated cells. Each point represents data from an individual culture, and all values are shown as mean ± SEM \*\*\**P* < 0.001, calculated by unpaired Student T test. **B)** Quantification of mouse DGKκ mRNA by qRT-PCR in RNA extracts of Cos-1 cells transfected with plasmid pCI-mDGKκ-HA and pre-transfected with siRNA control (siC) or siRNA against FMRP (siFMR1) or mock transfected (NT). Data are means of fold change ± SEM, determined using ΔΔCt method with Actb as normalizer, n = 3 biological replicates.

**Supplementary Fig. 2:**
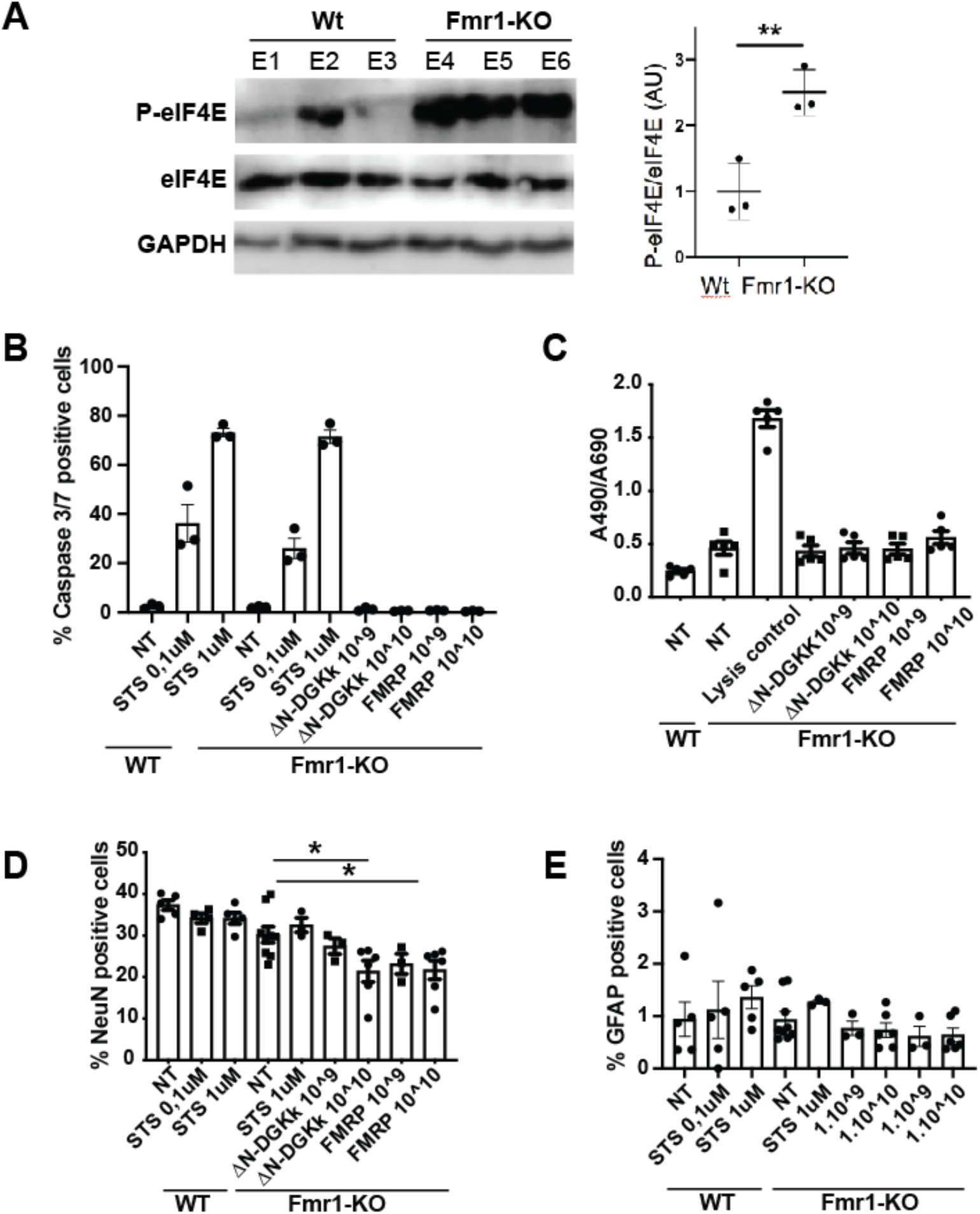
Phosphorylation of eIF4E is increased in Fmr1-KO cortical neurons compared to WT, and ΔN-DGKk expression in neurons does not lead to toxicity. **A)** Representative immunoblots of lysates from WT and Fmr1-KO cortical neurons and quantification of phosphorylation and total levels of eIF4E. GAPDH was used as a loading control. For quantification, the phospho-protein signal was normalized first against total protein signal and is presented relative to the signal for WT culture. Each point represents data from an individual culture, and all values are shown as mean ± SEM \*\**P* < 0.01 by Student T test (*n* = 3 individual cultures). Quantification of caspase 3/7 activity (**B**), release of lactate dehydogenase (LDH) (**C**), percentage of NeunN (**D**) and GFAP (**E**) positive cells, in WT and Fmr1-KO cortical neurons untreated (NT) or transduced at 8 DIV for 8 DIV with indicated titers of AAV (viral genome copies) AAVRh10-ΔN-DGKk or AAVRh10-FMRP by immunofluorescence high throughput cell imaging. Positive control wells were treated with apoptotic inducer staurosporine (STS) at 0.1 and 1μM for 6 hours. Data are mean ± SEM and analyzed using one-way ANOVA and Tukey’s multiple comparisons test. * p<0.05.

**Supplementary Fig. 3:**
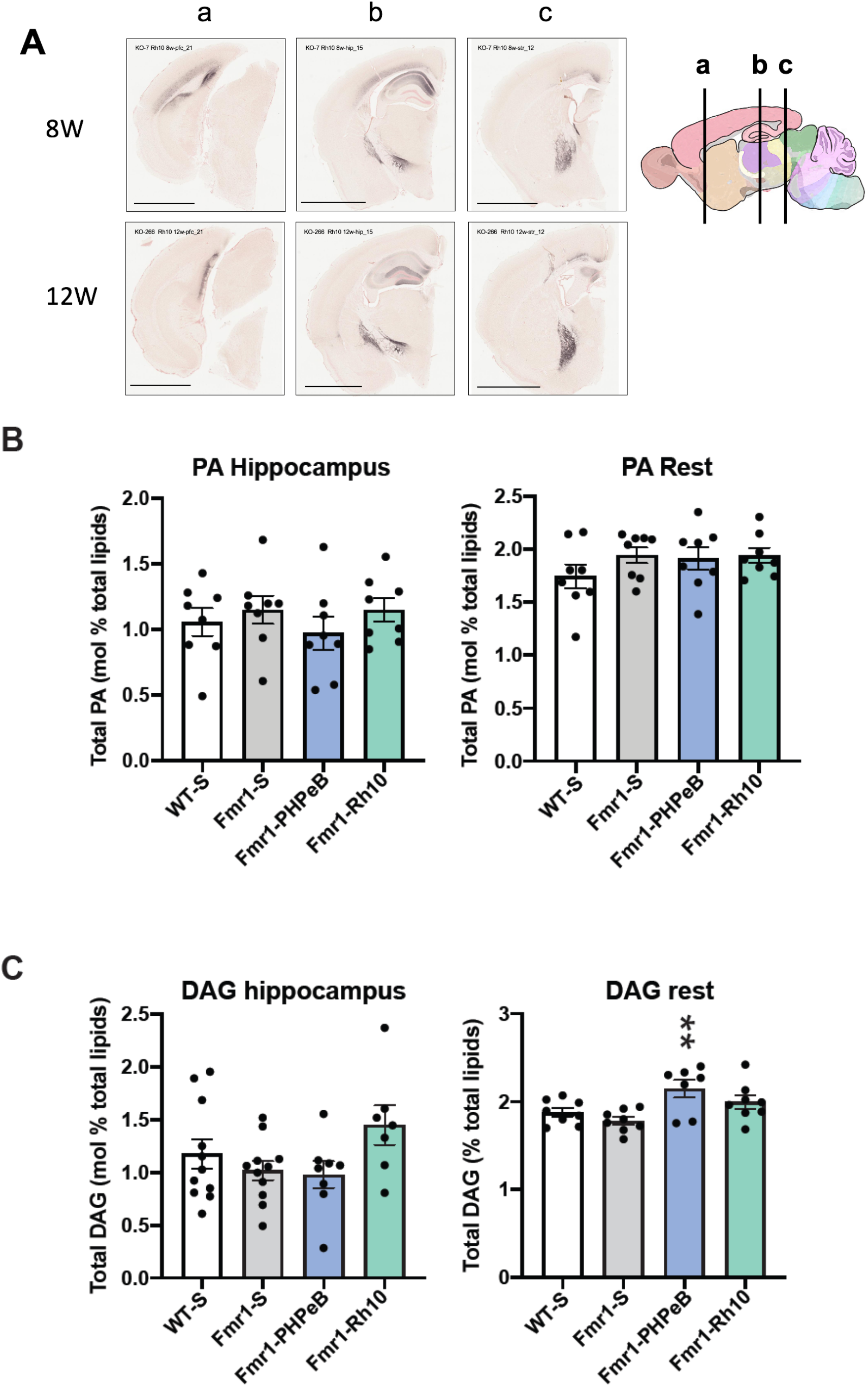
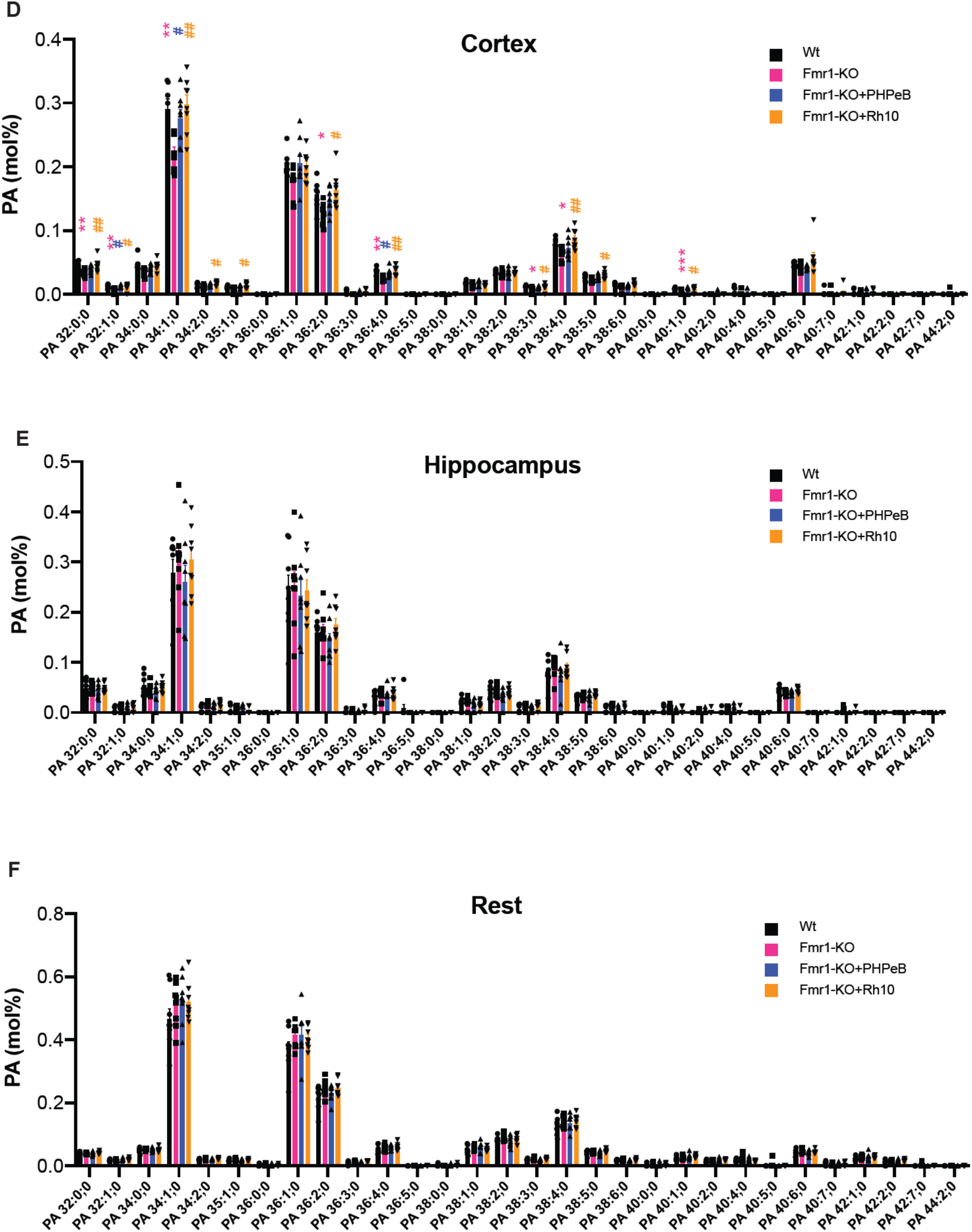
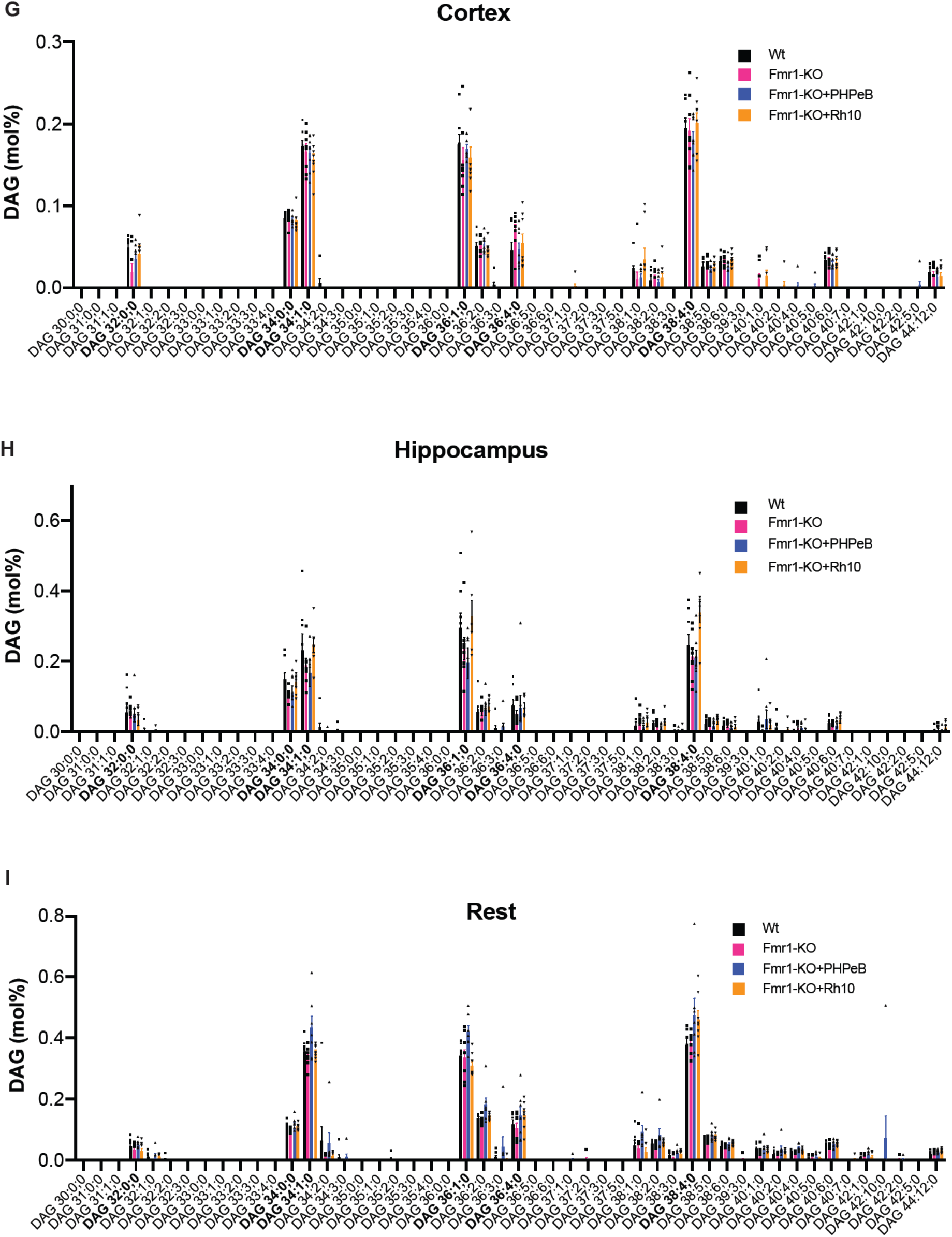
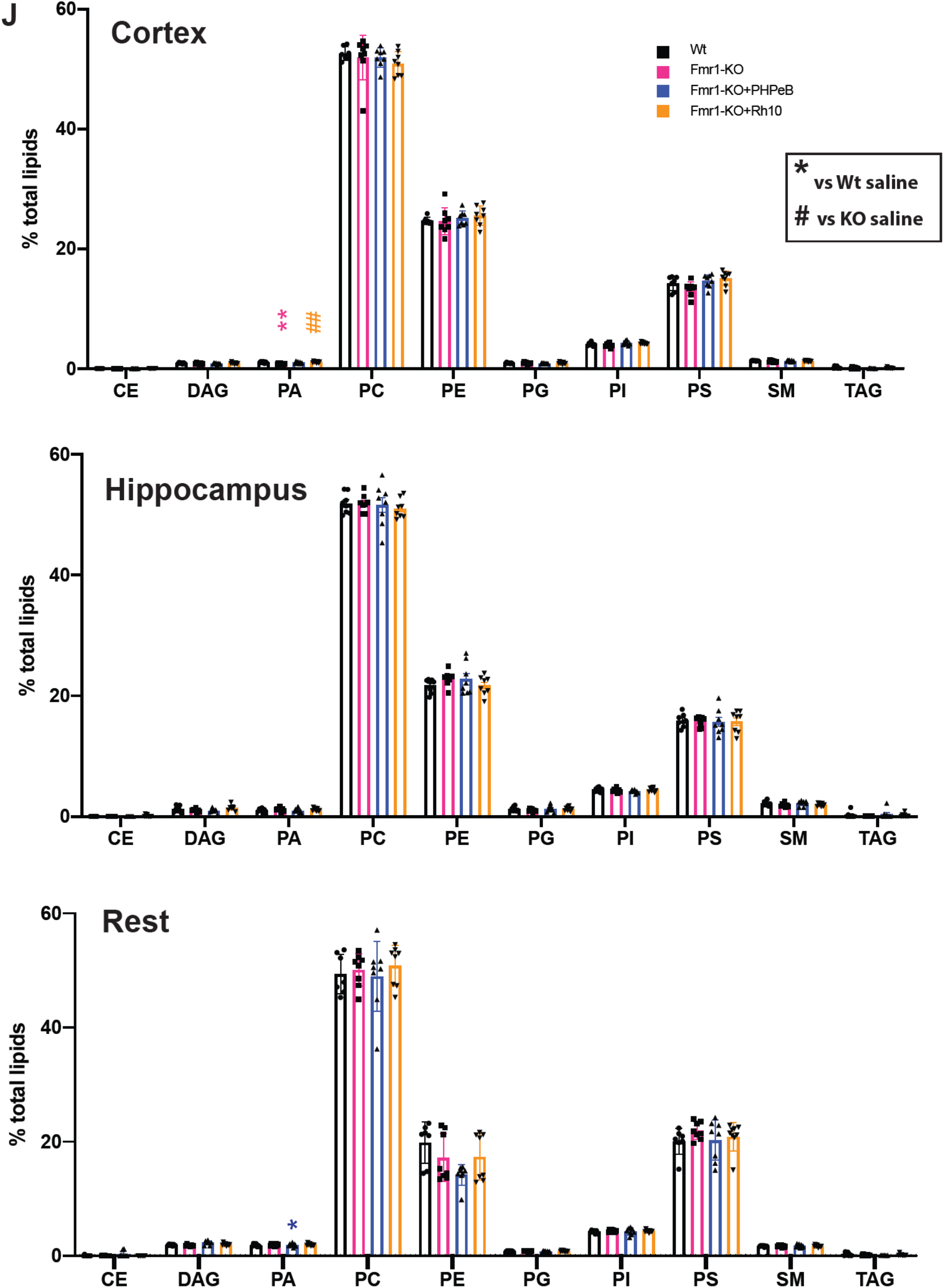
ΔN-DGKk expression in brain with AVV vectors is stable over time and normalizes phosphatidic acid level. **A)** Representative coronal brain sections processed for detection of ΔN-DGKk at 8 and 12 weeks post-injections using immunohistochemistry on Fmr1-KO mice treated with indicated treatment, counter stained with eosin hematoxylin. Three regions a, b, c, are shown with their corresponding position on brain sagital map. Scale bar 2mm. **B)** Measure of total phosphatidic acid (PA) level by mass spectrometry in hippocampus and rest of brain of WT mice treated with saline solution (WT) and Fmr1-KO mice treated with saline (Fmr1-S), AAVPHP.eB-ΔN-DGKk (Fmr1-PHP.eB), AAVRh10-ΔN-DGKk (Fmr1-Rh10) 8 weeks after injections. Data are expressed as mean ± SEM of mol % of total lipids and analyzed using one-way ANOVA and Tukey’s multiple comparisons test, *p<0.05, **p<0.01, n=8 individual animals, except for WT n=7. **C)** Total diacylglycerol (DAG) level in cortex measured as in B). PA and DAG individual species level respectively in cortex (**D, G**), hippocampus (**E, H**), and rest of brain (**F, I**). **J)** lipid composition (mol % of total lipid) for cholesterol esters (CE), diacylglycerol (DAG), phosphatidic acid (PA), phosphatidylcholine (PC), phosphatidylethanolamine (PE), phosphatidylglycerol (PG), phosphatidylinositol (PI), phosphatidylserine (PS), sphingomyelin (SM), triacylglycerol (TAG), measured as in **A**. Data are expressed as mean ± SEM and analyzed using two-way ANOVA. *p<0.05, **p<0.01, ***p<0.001, ****p<0.0001 vs WT-S, #p<0.05, ##p<0.01, ###p<0.001, ####p<0.0001 vs Fmr1-S.

**Supplementary Fig. 4:**
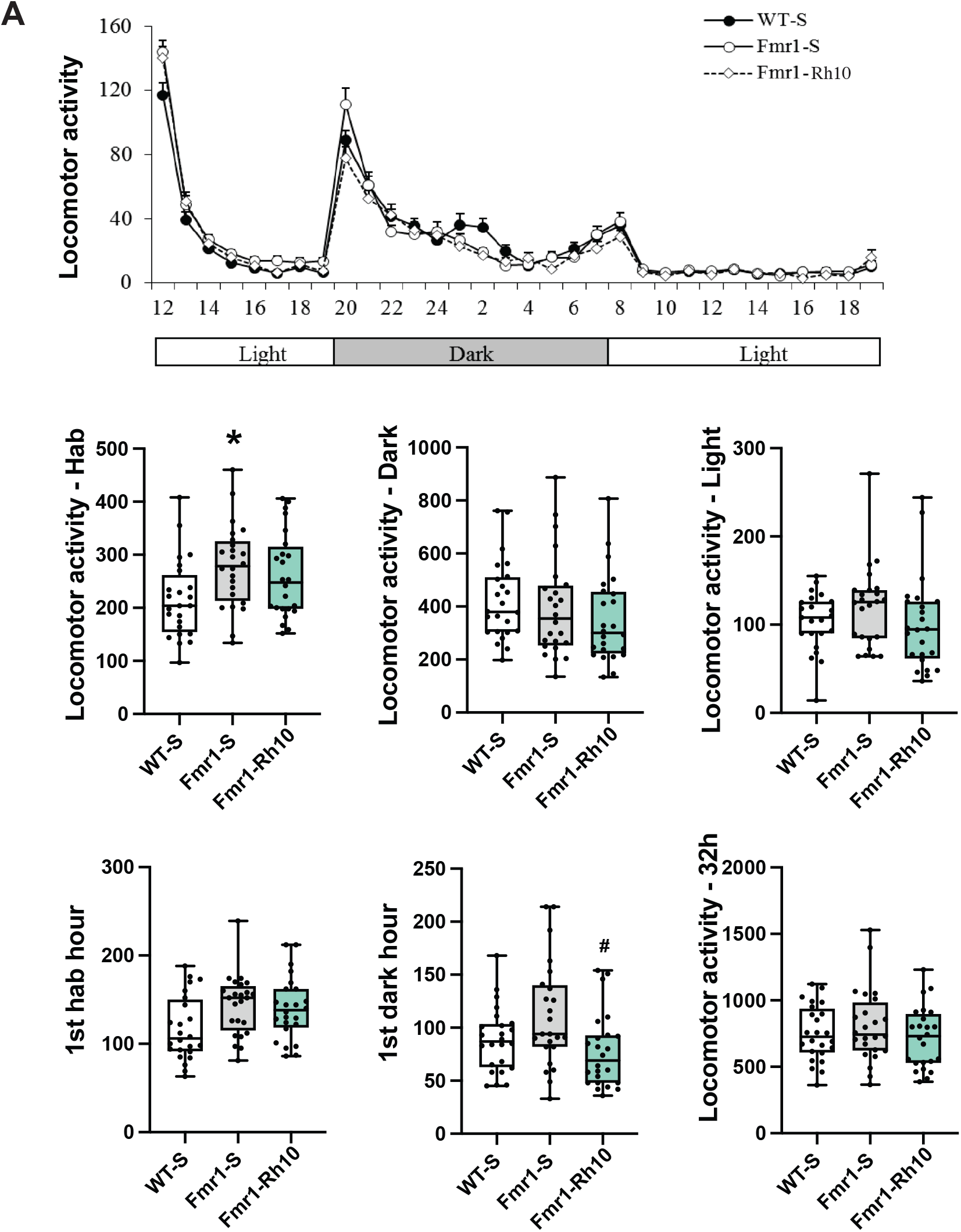
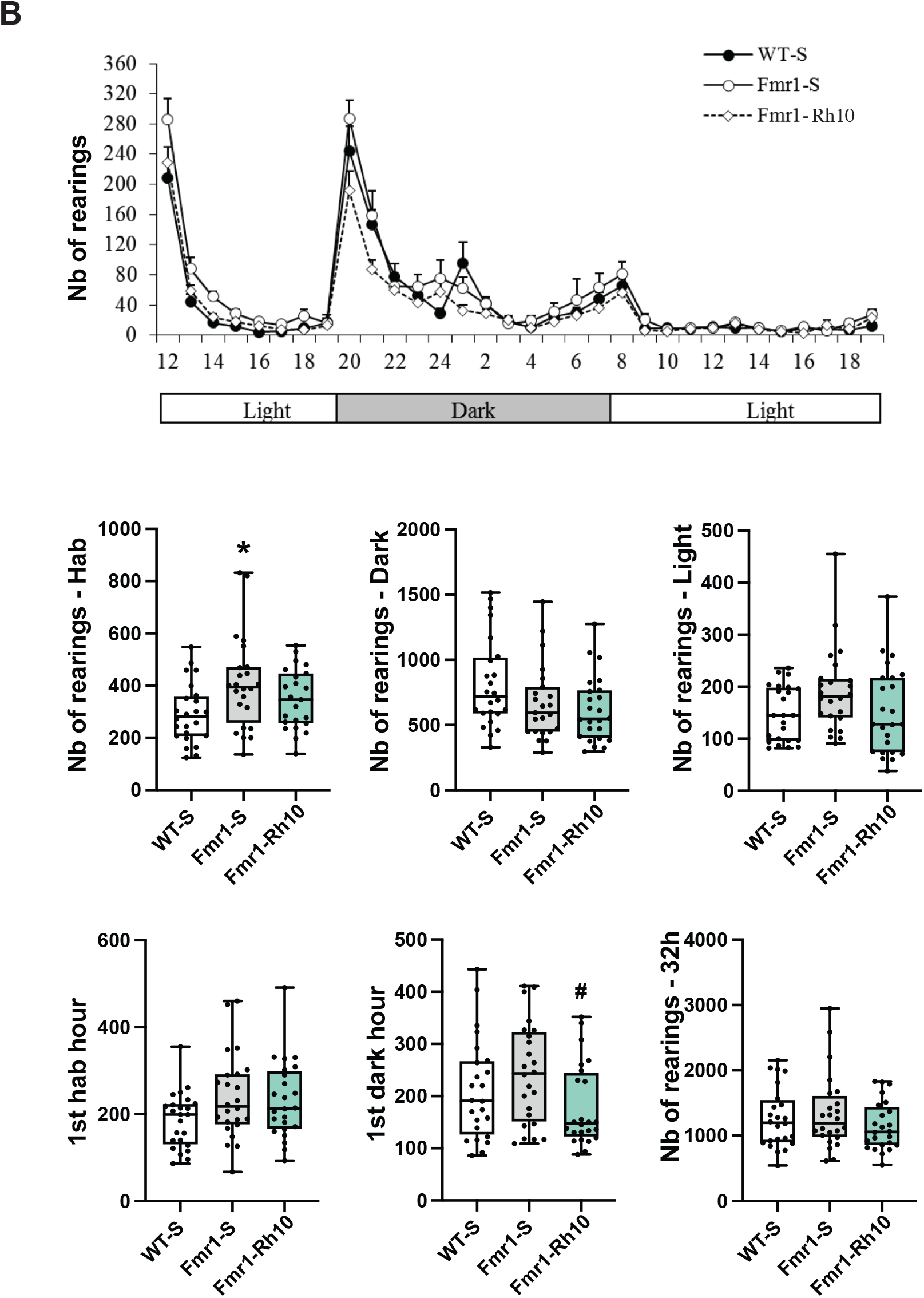
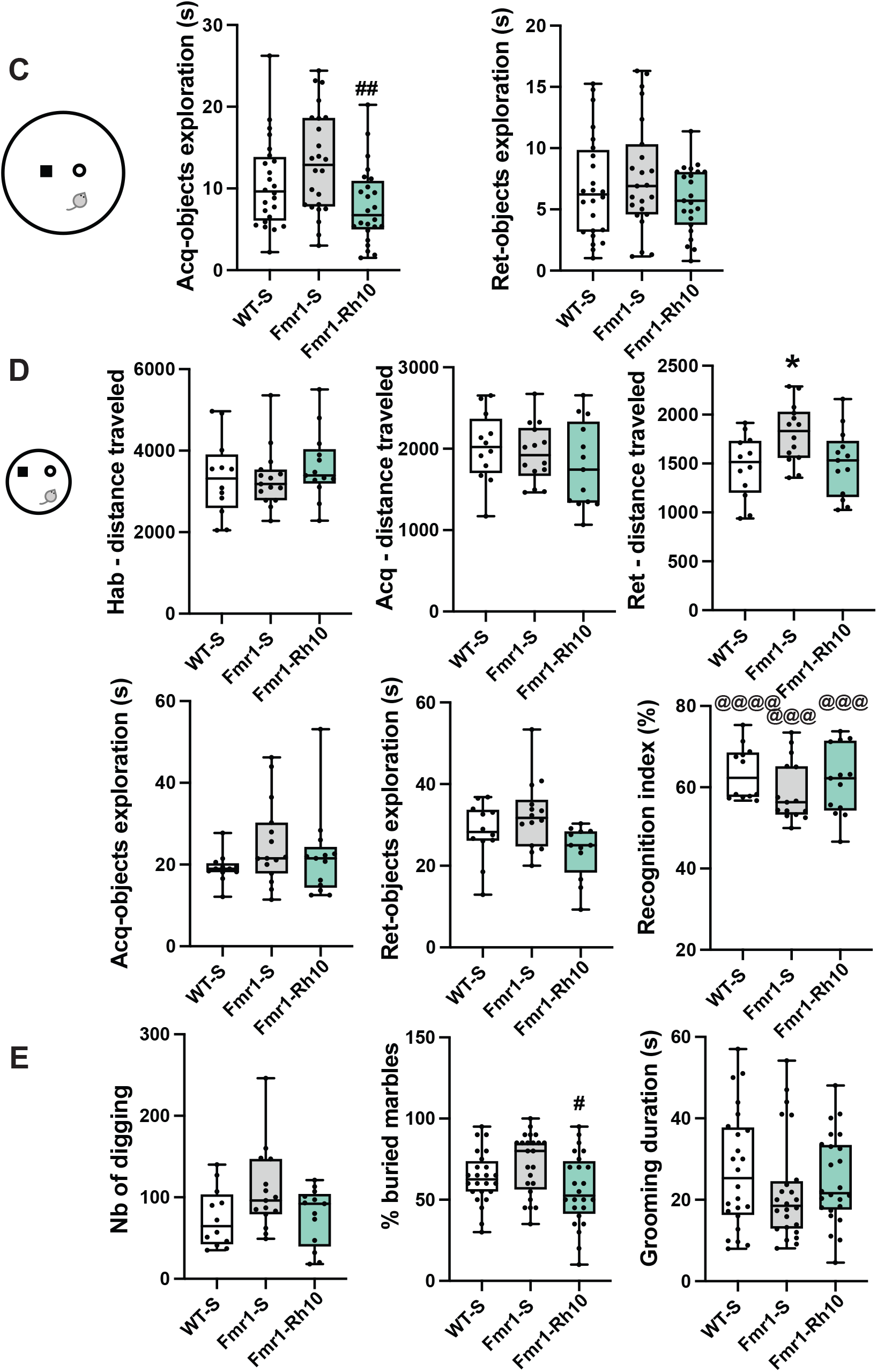
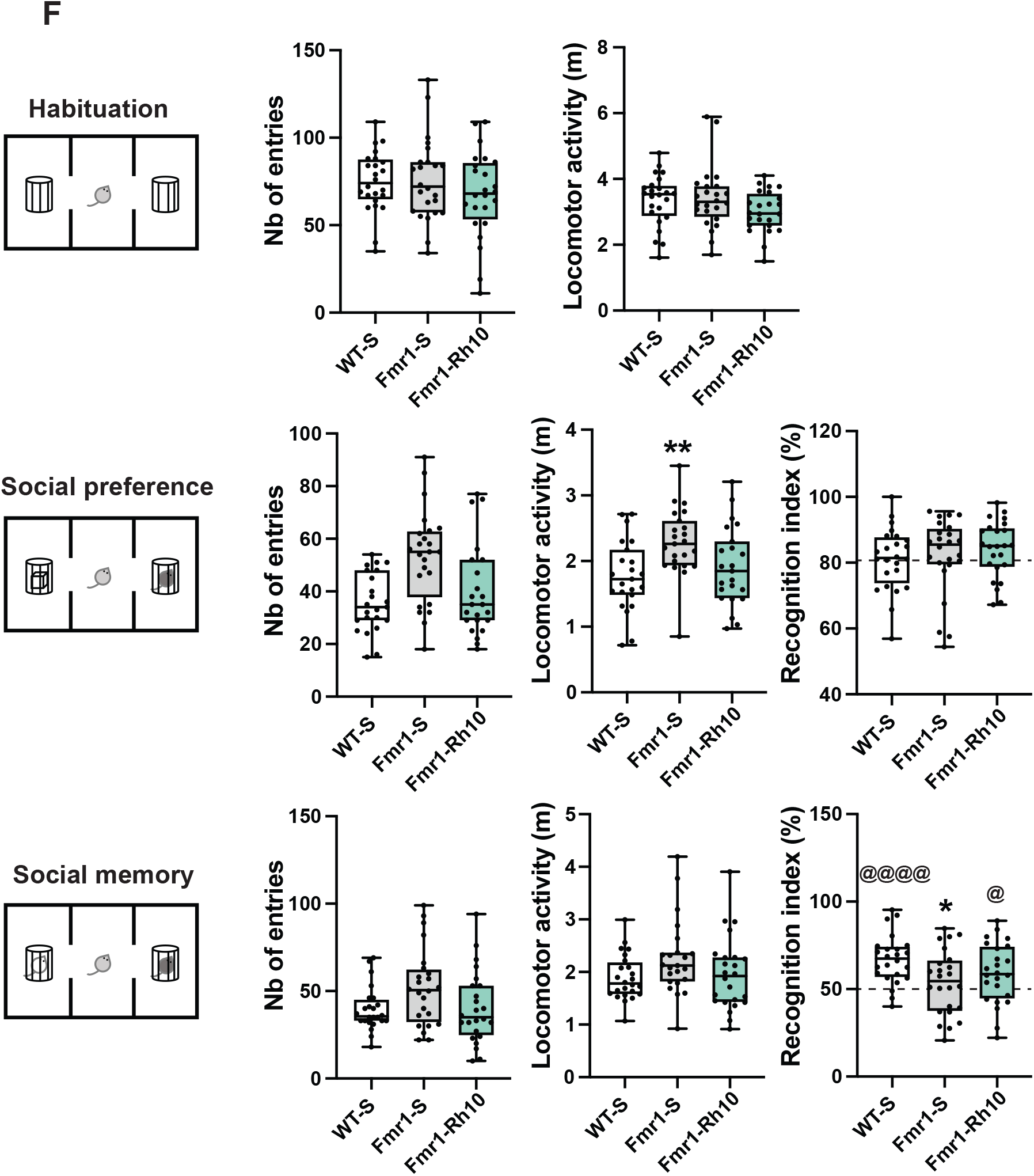
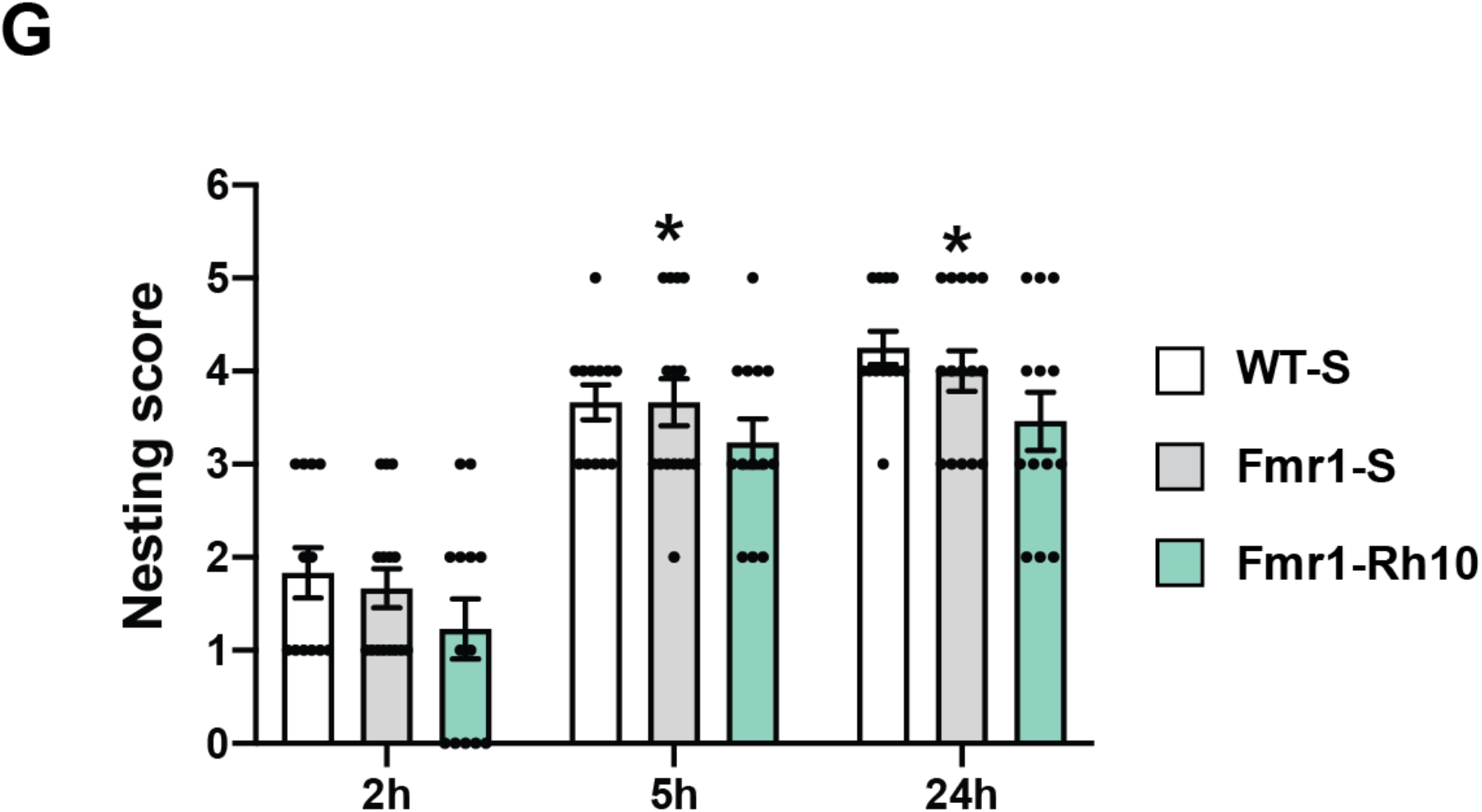
Behavioral analyses of AAVRh10-ΔN-DGKk treated *Fmr1*-KO mice (4 weeks after injection). **A)** Circadian activity (locomotor activity). Evolution of locomotor activity per hour over the 32h testing (upper panel) and total locomotor activity for the habituation, dark and light phases (mid panel) and for the first habituation hour, first dark hour and total duration (lower panel). **B)** Circadian activity (rearing activity). Evolution of rearing activity per hour over the 32h testing (upper panel) and total rearing activity for the habituation, dark and light phases (mid panel) and for the first habituation hour, first dark hour and total duration (lower panel). **C)** Novel object recognition in 50cm diameter arena (30cm height). Duration of objects exploration during the acquisition and retention trials. **D)** Novel object recognition in 30cm diameter arena (30cm height). Locomotor activity (distance) in the whole arena during the 15min habituation, acquisition and retention trials. Duration of objects exploration during the acquisition and retention trials and recognition index. **E)** Digging, marble burying and grooming duration tests. **F)** Social recognition. Number of entries and locomotor activity (total traveled distance in cm) in the two side compartments during habituation (up), social preference (middle) and social memory (bottom) sessions. Social preference was determined as percentage of exploration of a congener vs an object (middle right) and social memory as percentage of exploration of a novel vs familiar congener (bottom right). **G)** Nest building. Scoring of nests at 2, 5 and 24h. 0-5 scale as described by Gaskill et al (2013): 0 = undisturbed nesting material; 1 = disturbed nesting material but no nest site; 2 = a flat nest without walls; 3 = a cup nest with a wall less than ½ the height of a dome that would cover a mouse; 4 = an incomplete dome with a wall ½ the height of a dome; 5 = a complete dome with walls taller than ½ the height of a dome, which may or may not fully enclose the nest. Data are expressed as median with interquartile range with minimum and maximum values for A-F, mean ± SEM for G, and analyzed using one-way ANOVA and Tukey’s multiple comparisons test. * p<0.05 vs WT-S; # p<0.05 vs Fmr1-S; one group t-test. *p<0.05 vs WT-S, @@@ p<0.001, @@@@ p<0.0001 vs chance (50%); and χ2 test. *p<0.05 vs WT-S.

**Supplementary Fig. 5:**
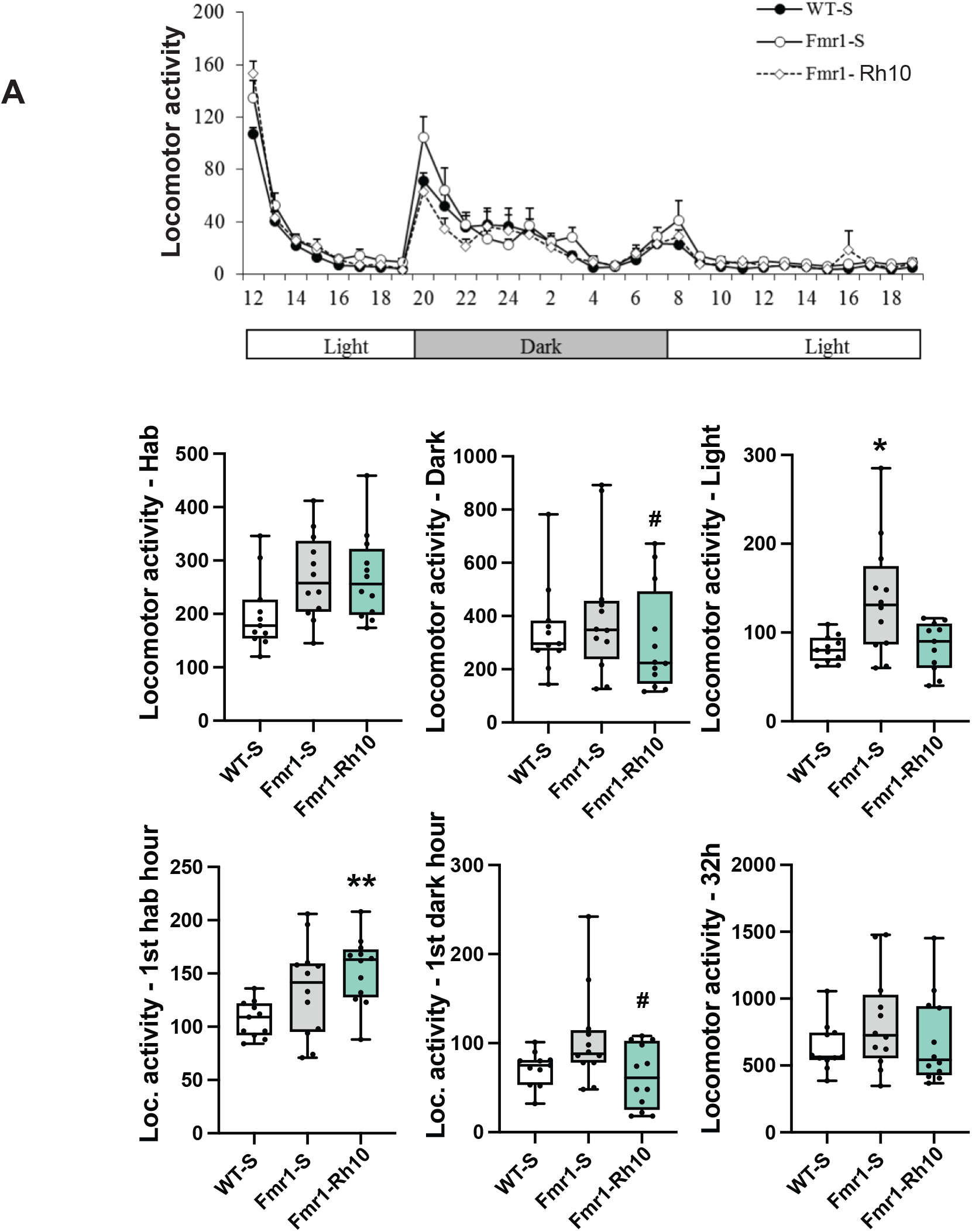
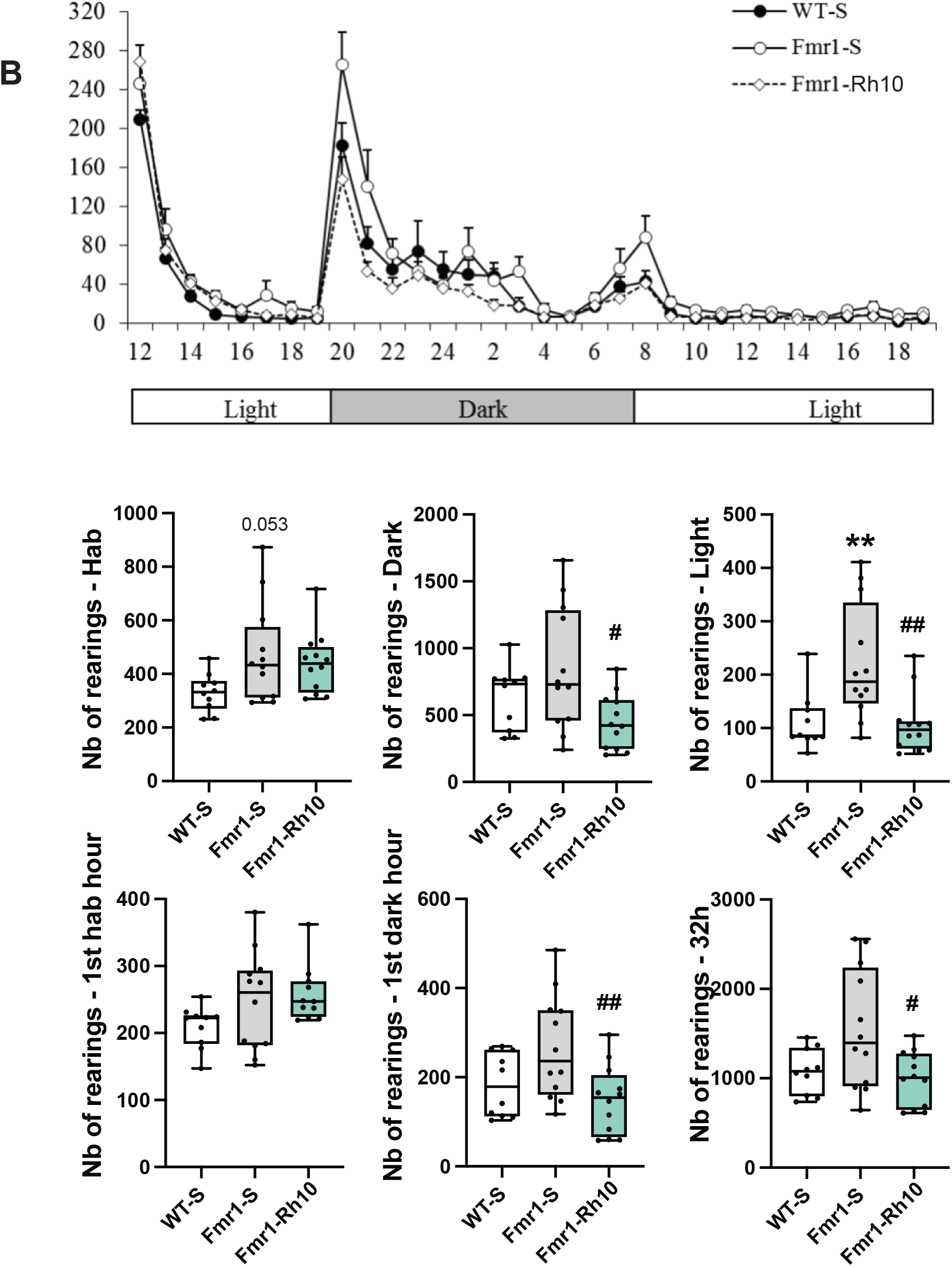
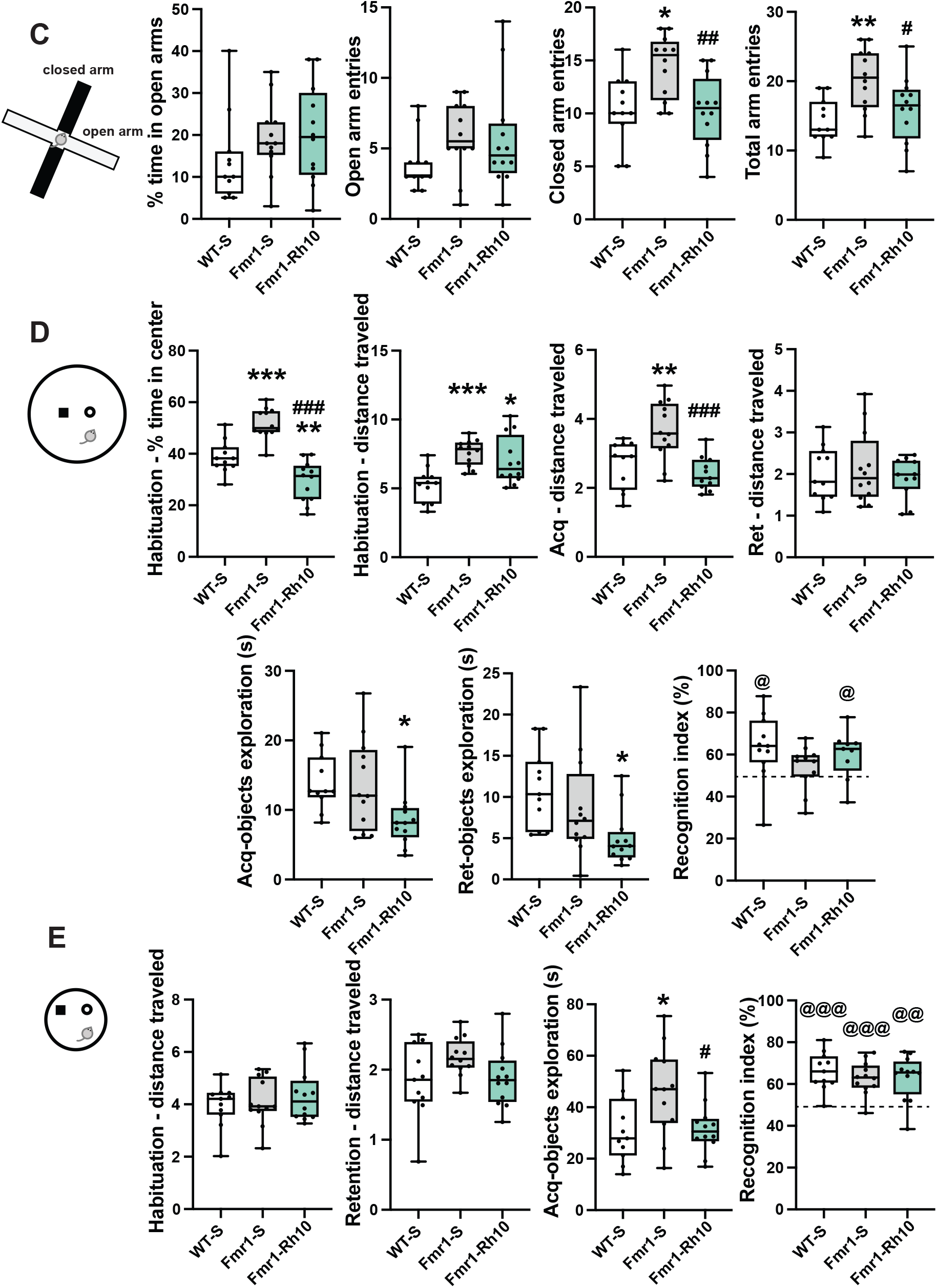
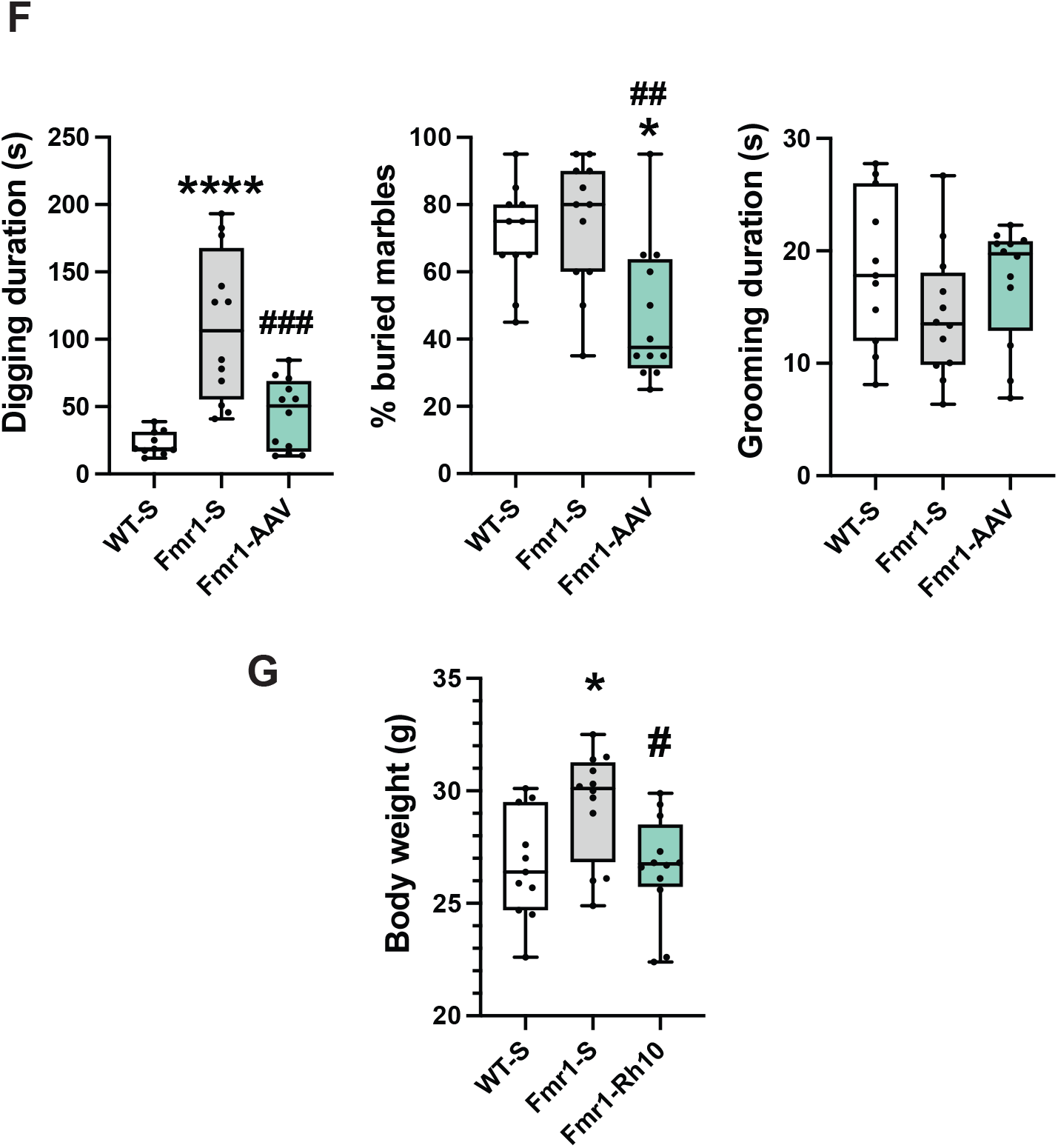
Behavioral analyses of AAVRh10-ΔN-DGKk treated *Fmr1*-KO mice (8 weeks after injection). Circadian activity analysis of locomotor (**A**) and rearing (**B**) activity per hour over the 32h testing (upper panel) and total locomotor and rearing activity for the habituation, dark and light phases (mid panel) and for the first habituation hour, first dark hour and total duration (lower panel). **C**) Elevated Plus Maze. Percentage of time spent in open arms and number of entries in open, closed and total (open+closed) arms. **D**) Novel object recognition in 50cm diameter arena (30cm height). Percentage of time spent in the center during the habituation, locomotor activity (distance) in the whole arena during the habituation, acquisition and retention trials. Duration of objects exploration during the acquisition and retention trials and recognition index. **E**) Novel object recognition in 30cm diameter arena (30cm height). Locomotor activity (distance) in the whole arena during the habituation and retention trials. Duration of objects exploration during the acquisition trials and recognition index. **F**) Digging, marble burying and grooming duration tests. **G**) Body weight of mice. Data are expressed as median with interquartile range with minimum and maximum values and analyzed using one-way ANOVA, Tukey’s multiple comparisons test and one group t-test. *p<0.05, **p<0.01, ***p<0.001 vs WT-S, # p<0.05, ## p<0.01, ### p<0.001 vs Fmr1-S, @p<0.05, @@p<0.01, @@@ p<0.001 vs chance (50%).

**Supplementary Fig. 6:**
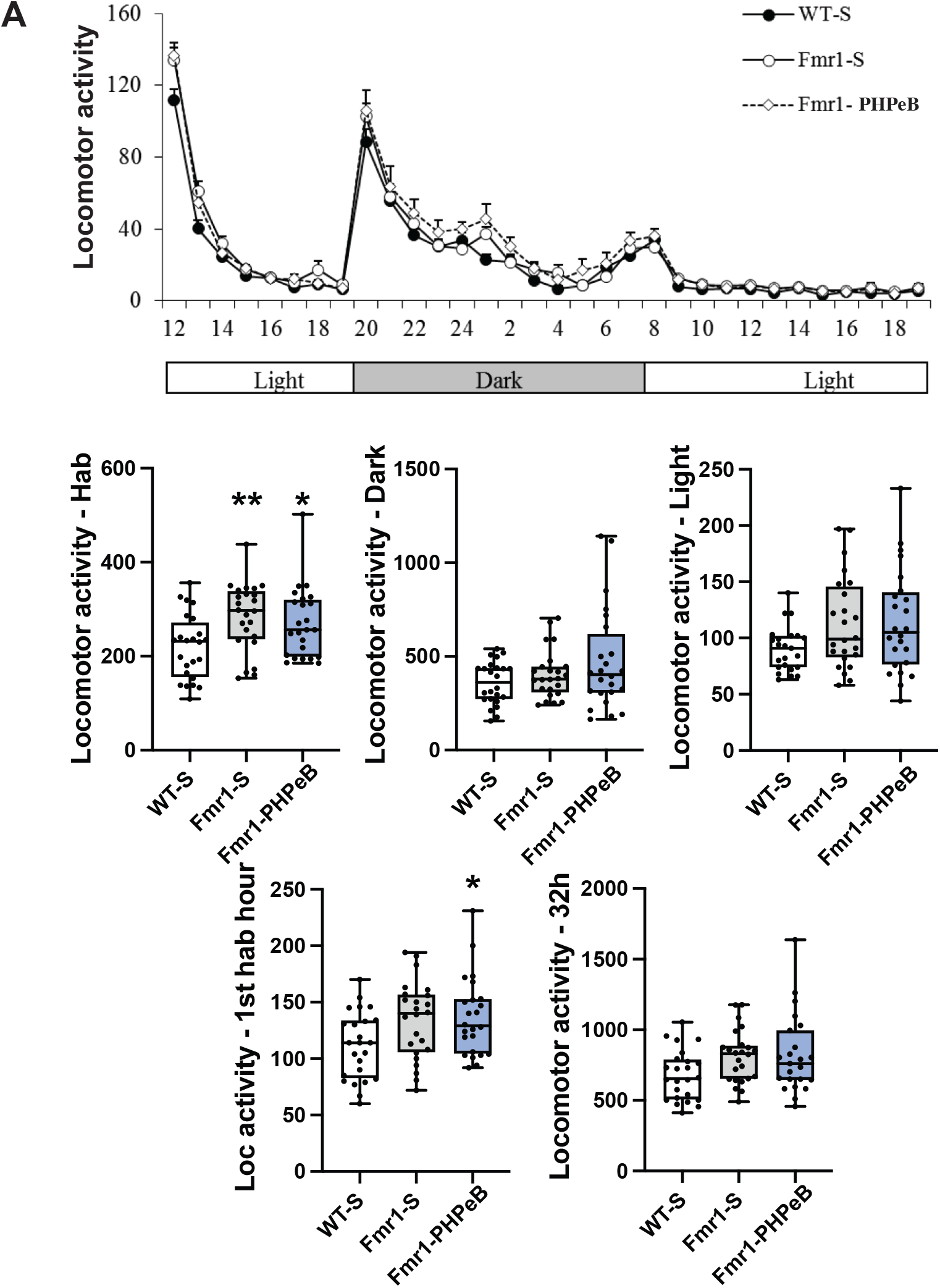
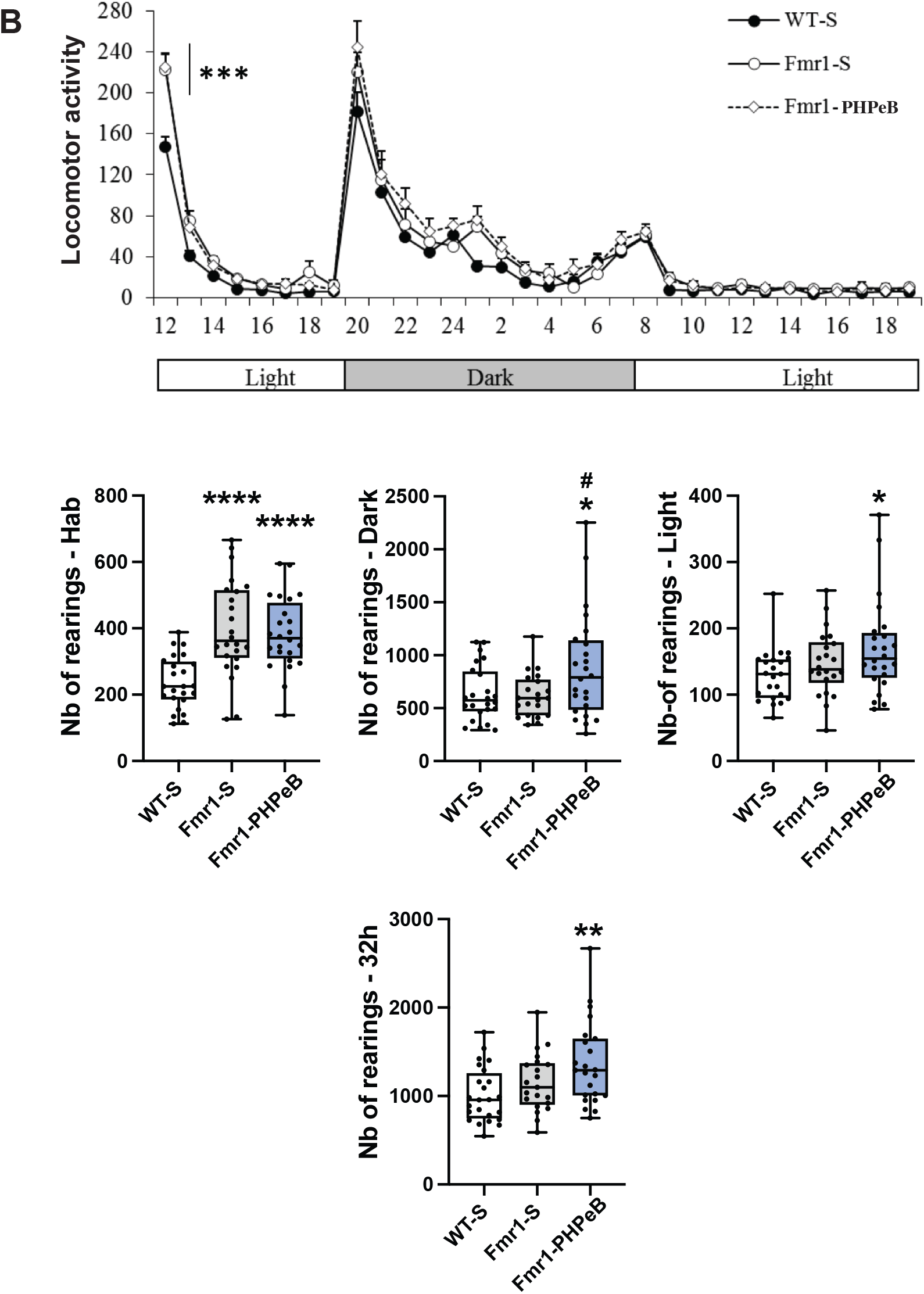
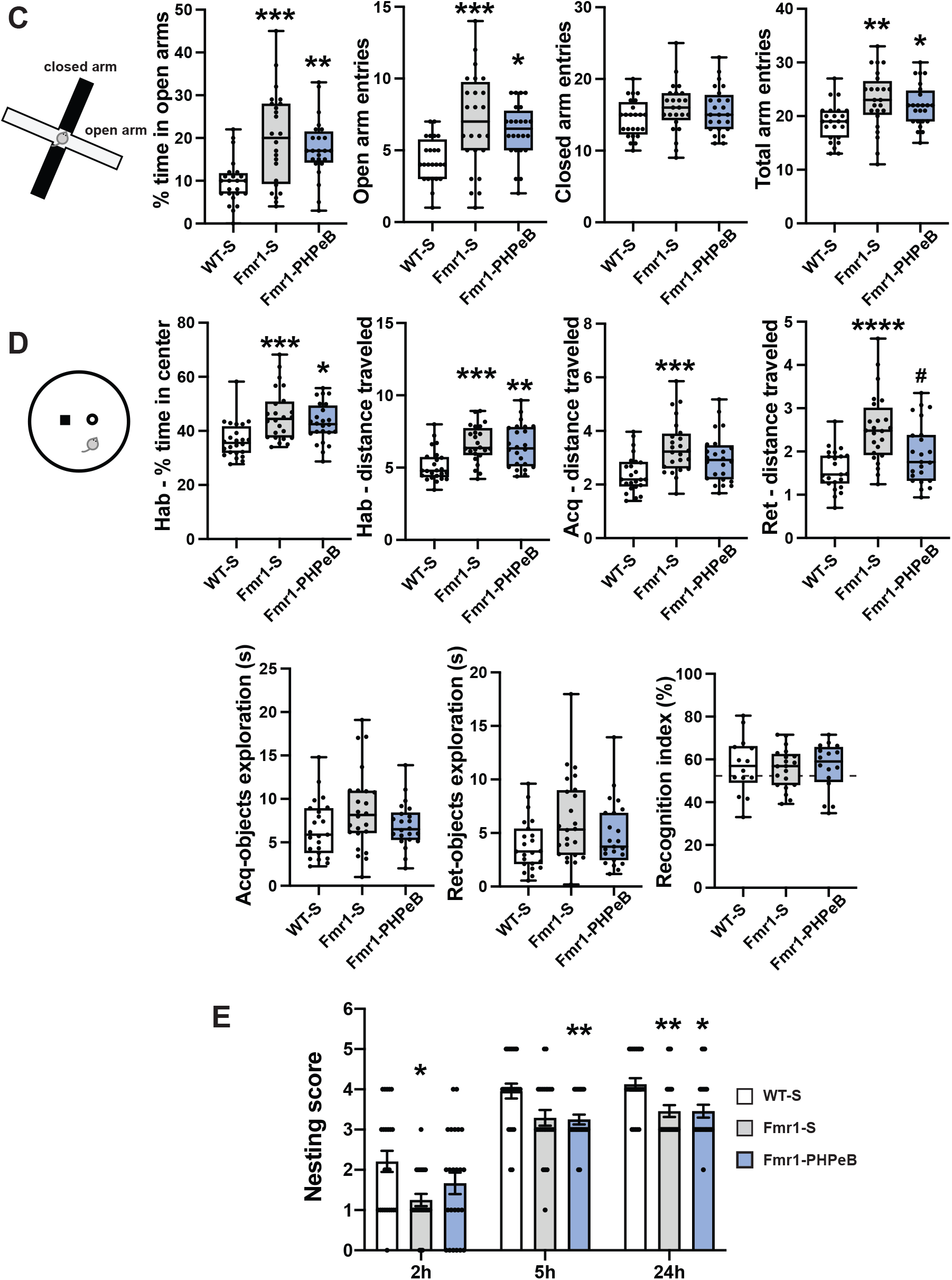
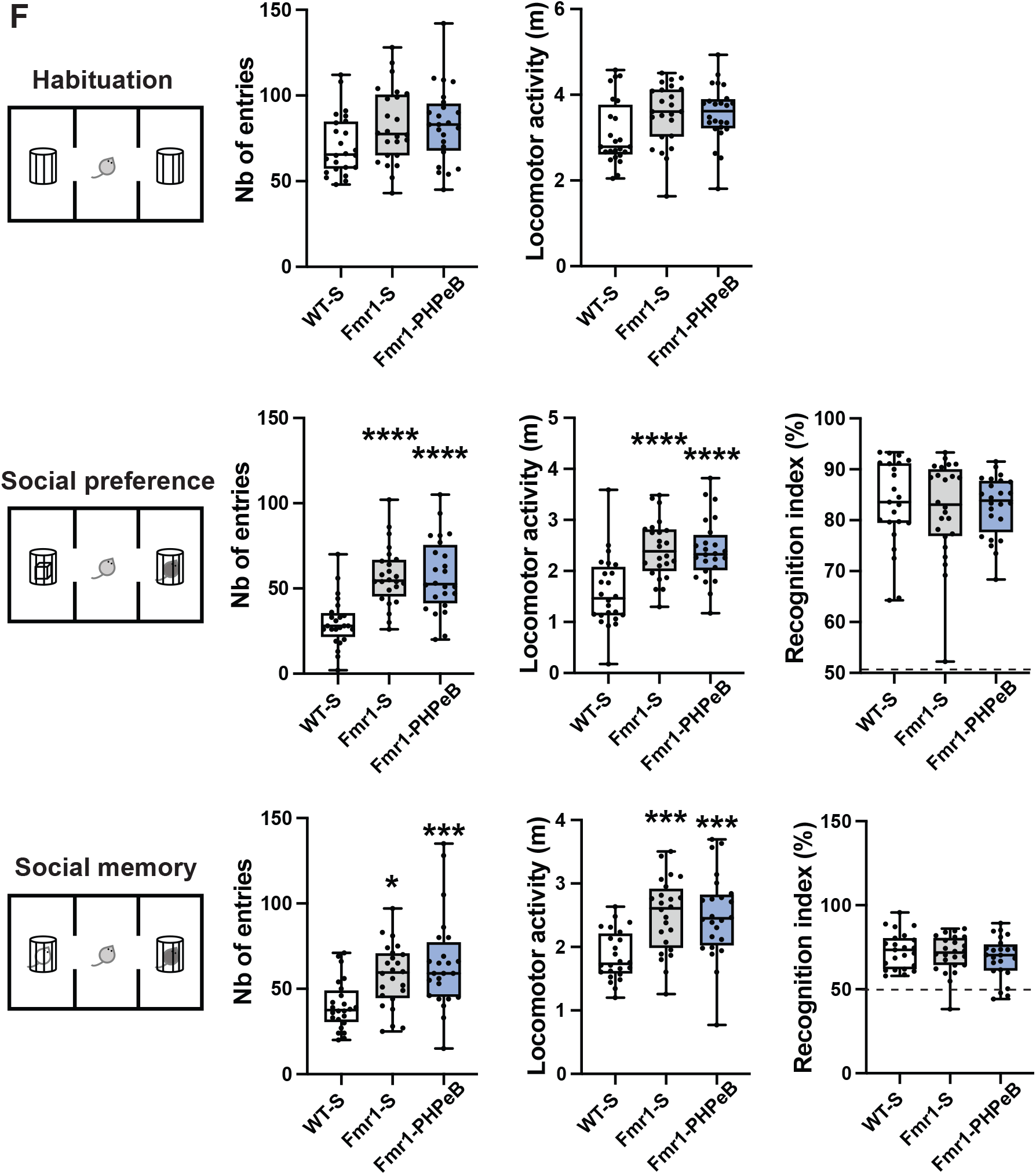
Behavioral analyses of AAVPHP.eB-ΔN-DGKk treated *Fmr1*-KO mice (4 weeks after injection). Circadian activity analysis of locomotor (**A**) and rearing (**B**) activity per hour over the 32h testing (upper panel) and total locomotor and rearing activity for the habituation, dark and light phases (mid panel) and for the first habituation hour, first dark hour and total duration (lower panel). **C**) Elevated Plus Maze. Percentage of time spent in open arms and number of entries in open, closed and total (open+closed) arms. Novel object recognition in 50cm diameter arena (30cm height). Percentage of time spent in the center during the habituation, locomotor activity (distance) in the whole arena during the habituation, acquisition and retention trials. Duration of objects exploration during the acquisition and retention trials and recognition index. **E)** Nest building. Scoring of nests at 2, 5 and 24h as in Sup. Fig. 4. **F)** Social recognition. Number of entries and locomotor activity in the two side compartments during habituation (up), social preference (middle), social memory (bottom) sessions. Social preference was determined as percentage of exploration of a congener vs an object (middle right) and social memory as percentage of exploration of a novel vs familiar congener (bottom right). Data are expressed as median with interquartile range with minimum and maximum values for A-F, as mean ± SEM for E and analyzed using one-way ANOVA, Tukey’s multiple comparisons test and one group t-test (recognition index) or χ2 test (nesting). *p<0.05 vs WT-S. *p<0.05, **p<0.01, ***p<0.001, ****p<0.0001 vs WT-S, # p<0.05 vs Fmr1-S.

**Supplementary Fig. 7.**
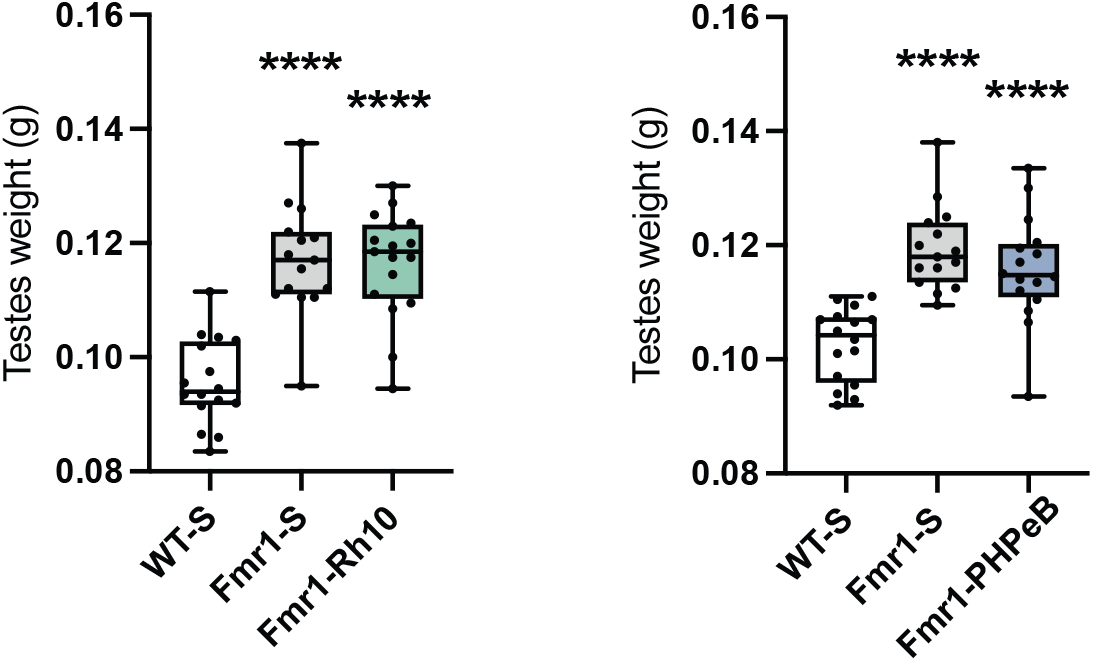
Testes weight of AAVRh10-ΔN-DGKk (left) and AAVPHP.eB-ΔN-DGKk (right) treated *Fmr1*-KO mice (12 weeks after injection). Data are means of the two testes of each animal and expressed as median with interquartile range with minimum and maximum values and analyzed using one-way ANOVA, Tukey’s multiple comparisons test ***p<0.0001 vs WT-S.

**Supplementary table 1:** Phenotyping pipeline.

| Age (weeks): | 5 | 6-8 | 9 | 10 | 11 | 12 | 13 | 14 | 15 | 16 |
| --- | --- | --- | --- | --- | --- | --- | --- | --- | --- | --- |
| Group 4W | Injection<br>AAVRh10<br>PHPeB | Monitoring<br>/acclimatation | Circadian<br>activity<br>Plus Maze | Open field<br>Grooming | Nesting<br>NOR<br>Marble<br>burying | Social<br>Recognition |  |  |  |  |
| Group 8W | Injection<br>AAVRh10<br>PHPeB | Monitoring<br>/acclimatation |  |  |  |  | Circadian<br>activity<br>Plus Maze | Open field<br>Grooming | Nesting<br>NOR<br>Marble<br>burying | Social<br>Recognition |

**Supplementary table 2:**
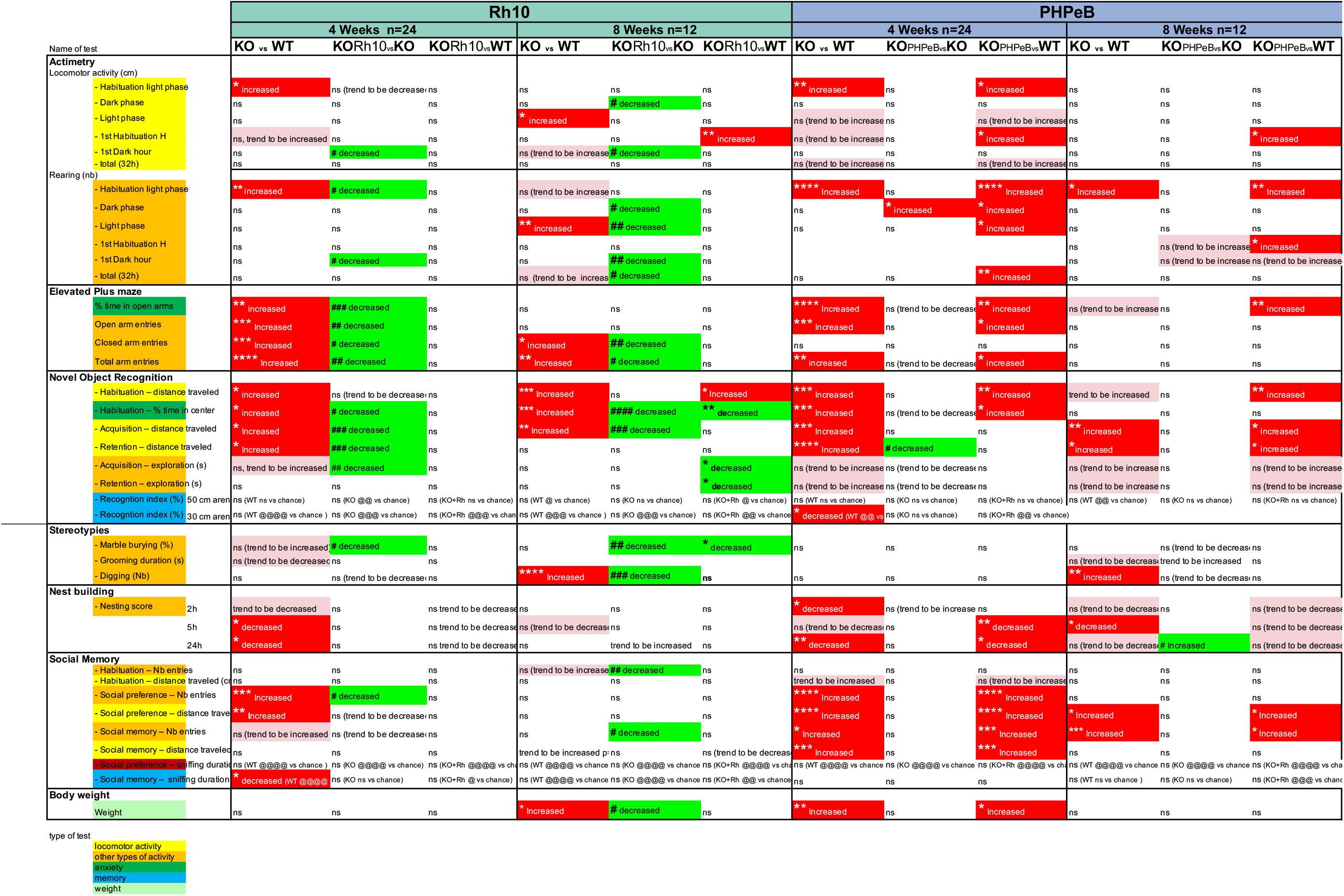
Summary of the phenotypic comparisons between WT mice treated saline (WT-S), *Fmr1*-KO treated saline (KO-S) and *Fmr1*-KO treated AAV (KO-Rh10 or KO-PhPeB) 4 and 8 weeks post-treatment. *, **, ***, **** p<0.05, 0,01, 0,001, 0,0001 KO-S vs WT-S, or KO-AAV vs WT-S. #, ##, ##, #### p<0.05, 0,01, 0,001, 0,0001 KO-AAV vs KO. *p<0.05, **p<0.01, ***p<0.001, ****p<0.0001 vs WT-S, # p<0.05 vs Fmr1-S.

